# Described eukaryotic species in published species-level phylogenies: placement by molecular data, dates and date uncertainty

**DOI:** 10.64898/2026.05.31.729127

**Authors:** Francisco Richter

## Abstract

Comparative analyses take published species-level phylogenies as data. Their estimates depend on how many described species a tree holds, how many were placed by sequence data, and whether the tree is dated with an uncertainty that can be carried into the analysis. These properties are reported one tree at a time, not on a common basis across eukaryotes.

I divided the 2,059,063 described eukaryotic species into 60 groups and took the largest published species-level tree found for each of the 24 groups that have one as its reference tree. Resolving tips to Catalogue of Life accepted names, I counted species at four nested levels: in the tree, placed with molecular data, in a tree dated in absolute time, and in a dated tree whose archive carries date uncertainty (a set of dated trees or node intervals). I then fitted a constant-rate birth–death model to the 7 dated trees that hold taxonomically placed tips, pruned to their sequenced tips, with the sampling fraction that matches the pruned tree and with the full tree’s fraction.

The species-level trees found name 19.4% of described eukaryotic species, and the reference trees 19.2%; the reference tree of each group places 7.6% with molecular data and 5.5% in a dated tree. Date uncertainty is available for 1.2%, 38% of them in the tree of Birds; posterior or bootstrap datings of sequence data cover 0.65%. A tree the search did not find can only add to that 19.4%. Keeping the full tree’s fraction after pruning lowers the estimated speciation rate in every group, and puts the estimate on the boundary of zero extinction in 100 of 100 squamate and 89 of 100 amphibian posterior trees. A likelihood-ratio test of the full tree’s fraction is made only on the 3 binary summary trees whose node ages come from a dating analysis (Fish, Mammals and Sharks); it rejects on none of them.

A per-group table gives the largest tree found, its sequenced tips, what its archive carries for date uncertainty and the sampling fractions that match the full and the pruned tree. The sources of all 7 trees with taxonomically placed tips also distribute a tree of their sequenced species, 4 of them dated.

## Introduction

Comparative and macroevolutionary analyses take species-level phylogenies as data. Diversification rates, trait models, phylogenetic diversity and phylogenetic regressions are estimated on a tree that someone else built and archived, and the estimates depend on how many species the tree holds and on how those species were placed and dated (Rangel et al., 2015; Title and Rabosky, 2019). A species attached to a tree by taxonomy, without sequence data (a taxonomically placed tip), carries no information about its branch lengths, and diversification rates estimated on trees completed in this way are biased (Rabosky, 2015; Chang et al., 2020). The fraction of species a tree samples enters birth–death likelihoods directly (Stadler, 2009; FitzJohn et al., 2009), and ages taken as fixed point estimates understate the uncertainty of every quantity computed from them (Rangel et al., 2015; de Villemereuil et al., 2012).

The gap between described species and species in a phylogeny, the Darwinian shortfall (Hortal et al., 2015; Diniz-Filho et al., 2013), has been measured for named or sequenced species within single clades or regions (Rudbeck et al., 2022; Guedes et al., 2024). The Open Tree of Life combines published trees with a taxonomy and reports which of its tips are placed by an input phylogeny (Hinchliff et al., 2015); TimeTree and DateLife summarise divergence times from published studies (Kumar et al., 2022; Sánchez Reyes et al., 2024); TreeHub indexes published trees (Wu et al., 2025); and the VertLife portal (https://vertlife.org) distributes tree sets of birds, mammals, amphibians, squamates and sharks, including, for several of them, versions restricted to sequenced species (Jetz et al., 2012; Upham et al., 2019; Jetz and Pyron, 2018; Tonini et al., 2016; Stein et al., 2018). None of these reports, for every group of eukaryotes on a common basis, how many described species the most complete available tree places with molecular data, whether that tree is dated in absolute time, and whether its archive carries an uncertainty on the dates that an analysis can use.

Counting accepted species of one name authority over described species puts every group on the same footing. For a user, the count says how far a downloaded tree falls short of its group and why. Summed over groups, it shows where sequence data, dates and date uncertainty are missing from archived trees.

I count, for the largest published species-level tree found for each group of eukaryotes, the described species it names, places with molecular data, places in a chronogram in absolute time, and places in a chronogram whose archive carries variation in node dates that a user can propagate. The result is a table (Table 1) that a user can consult before fitting a model to a group’s tree: which tree to use, the sampling fraction that matches the full tree and the tree pruned to its sequenced tips, and whether the source archive carries a tree set or node intervals with which date uncertainty can be carried into the analysis. A birth–death model fitted to the 7 dated trees that contain taxonomically placed tips shows how far the estimates move when the sampling fraction does not match the tree.

**Table 1.**
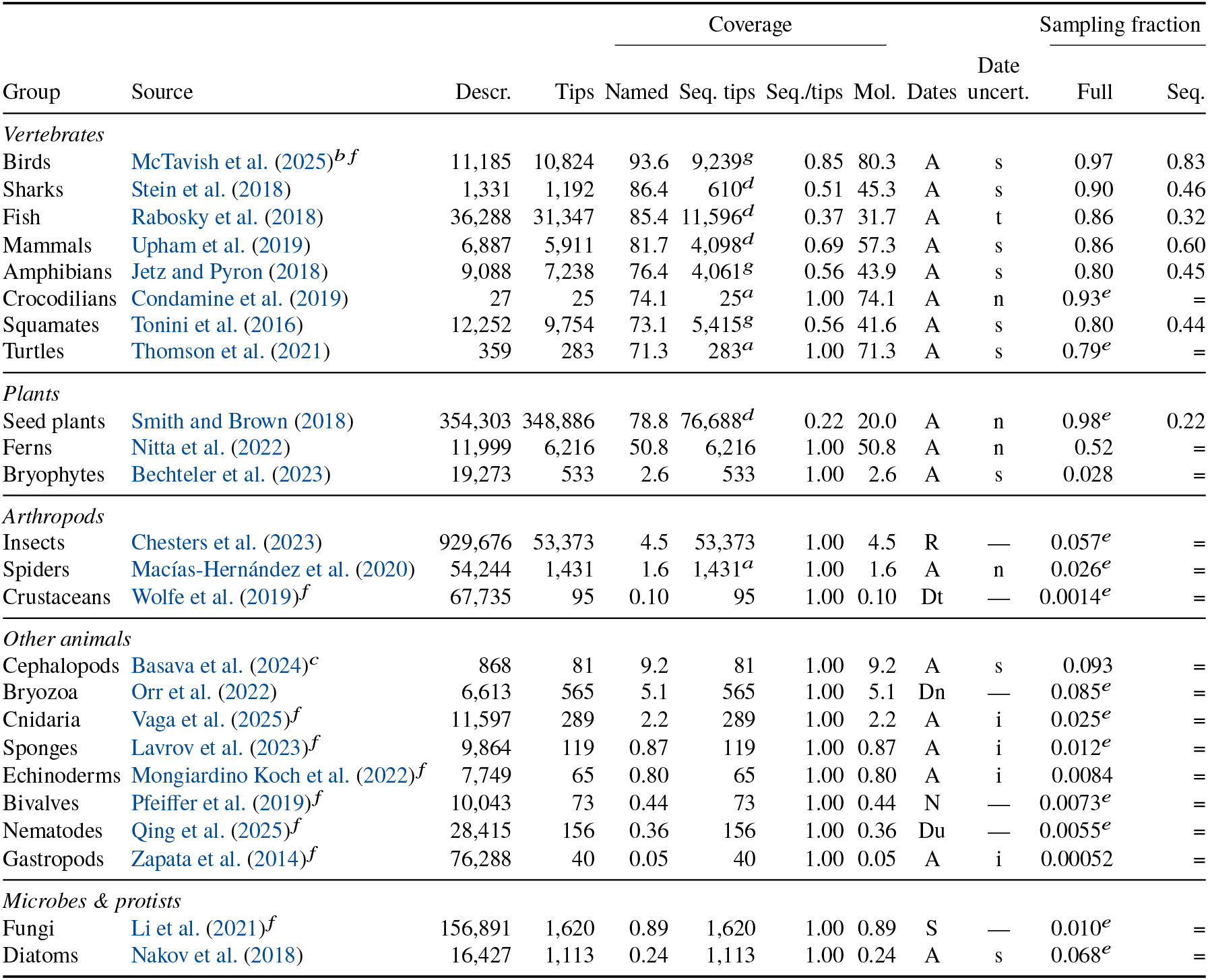
The reference tree of each eukaryotic group and what it supports. Descr.: described species of the group (the denominator). Tips: after removing tips outside the group, one per species. Named and Mol.: percentage of described species named in the tree, and named and placed with molecular data, counted as accepted species, hence below tips over described species where labels do not resolve or merge. Seq. tips: tips in the molecular analysis that built the tree (sequenced tips); Seq./tips: their share of the tips (the molecular fraction). Dates: A, chronogram in absolute time; R, ultrametric, relative time; for a tree that is not a chronogram, what the source study did: S, dated in relative time only; Dt, dated, archived chronogram lacks tips; Dn, dated, chronogram not archived; Du, dated, archive not readable; N, not dated. Date uncert., for a chronogram in absolute time, what the source archive carries that a user can propagate: s, a tree set whose node ages vary; i, intervals on the nodes of a summary tree; t, a tree set that varies only the placement of taxonomically added tips (no date uncertainty); n, nothing; Supplement S6 compares each archived crown interval with its calibrations. Sampling fraction: tips over described species, for fitting the full tree, and sequenced tips over described species, for fitting the tree pruned to its sequenced tips; both assume one described species per tip. An equals sign marks groups where every tip is sequenced, so the two fractions coincide. The trees of Bivalves, Cnidaria, Crustaceans, Echinoderms, Gastropods, Nematodes and Sponges sample one subclade or are phylogenomic backbones of a few dozen to a few hundred species; their rows describe the largest tree the search found, not how much of the group published trees cover (larger trees the search missed are marked *f*), and their fractions complete the row and are not inputs for fitting a model. *a*Sequenced tips stated by the source; the other groups’ sequenced tips are listed. *b*Automated synthesis of published trees and taxonomy; its sequenced tips are the tips of the source’s tree of tips an input phylogeny places (Materials and methods). *c*Source published as a preprint. *d*The source also distributes a dated tree of its sequenced species. *g*The source also distributes a tree of its sequenced species that is not dated (branch lengths in substitutions, or none). *e*Tips exceed the named species by more than 10% (Supplement S3). *f* A larger tree of the group or of a subclade, which the search did not find and which is not counted here, or, for Birds, a larger version of the reference tree’s synthesis aligned to a later taxonomy, not used by the rule stated in Materials and methods; the row describes the reference tree, and the size is as its source states it: Birds: McTavish et al. (2025), 11,017 tips of Aves in the Clements 2023 taxonomy (same AvesData 1.3 release), dated; Bivalves: Combosch et al. (2017), 219 ingroup species of Bivalvia; Cnidaria: Huang and Roy (2015), 1,547 species of Scleractinia, dated; Crustaceans: Copilaş-Ciocianu et al. (2020), 201 ingroup terminals of Amphipoda, dated and Wolfe et al. (2024), 344 tips of Brachyura, dated; Echinoderms: O’Hara et al. (2017), 576 species of Ophiuroidea, dated; Fungi: Varga et al. (2019), 5,284 species of Agaricomycetes, dated; Gastropods: Puillandre et al. (2014), 320 species of Conus; Nematodes: van Megen et al. (2009), 1,215 SSU rDNA sequences of Nematoda; Sponges: Redmond et al. (2013), more than 500 new 18S sequences of Demospongiae.

| Group | Source | Descr. | Tips | Coverage |  |  |  | Dates | Date uncert. | Sampling fraction |  |
|---|---|---|---|---|---|---|---|---|---|---|---|
|  |  |  |  | Named | Seq. tips | Seq./tips | Mol. |  |  | Full | Seq. |
| *Vertebrates* |  |  |  |  |  |  |  |  |  |  |  |
| Birds | McTavish et al. (2025)<sup>*b f*</sup> | 11,185 | 10,824 | 93.6 | 9,239<sup>*g*</sup> | 0.85 | 80.3 | A | s | 0.97 | 0.83 |
| Sharks | Stein et al. (2018) | 1,331 | 1,192 | 86.4 | 610<sup>*d*</sup> | 0.51 | 45.3 | A | s | 0.90 | 0.46 |
| Fish | Rabosky et al. (2018) | 36,288 | 31,347 | 85.4 | 11,596<sup>*d*</sup> | 0.37 | 31.7 | A | t | 0.86 | 0.32 |
| Mammals | Upham et al. (2019) | 6,887 | 5,911 | 81.7 | 4,098<sup>*d*</sup> | 0.69 | 57.3 | A | s | 0.86 | 0.60 |
| Amphibians | Jetz and Pyron (2018) | 9,088 | 7,238 | 76.4 | 4,061<sup>*g*</sup> | 0.56 | 43.9 | A | s | 0.80 | 0.45 |
| Crocodilians | Condamine et al. (2019) | 27 | 25 | 74.1 | 25<sup>*a*</sup> | 1.00 | 74.1 | A | n | 0.93<sup>*e*</sup> | = |
| Squamates | Tonini et al. (2016) | 12,252 | 9,754 | 73.1 | 5,415<sup>*g*</sup> | 0.56 | 41.6 | A | s | 0.80 | 0.44 |
| Turtles | Thomson et al. (2021) | 359 | 283 | 71.3 | 283<sup>*a*</sup> | 1.00 | 71.3 | A | s | 0.79<sup>*e*</sup> | = |
| *Plants* |  |  |  |  |  |  |  |  |  |  |  |
| Seed plants | Smith and Brown (2018) | 354,303 | 348,886 | 78.8 | 76,688<sup>*d*</sup> | 0.22 | 20.0 | A | n | 0.98<sup>*e*</sup> | 0.22 |
| Ferns | Nitta et al. (2022) | 11,999 | 6,216 | 50.8 | 6,216 | 1.00 | 50.8 | A | n | 0.52 | = |
| Bryophytes | Bechteler et al. (2023) | 19,273 | 533 | 2.6 | 533 | 1.00 | 2.6 | A | s | 0.028 | = |
| *Arthropods* |  |  |  |  |  |  |  |  |  |  |  |
| Insects | Chesters et al. (2023) | 929,676 | 53,373 | 4.5 | 53,373 | 1.00 | 4.5 | R | — | 0.057<sup>*e*</sup> | = |
| Spiders | Macías-Hernández et al. (2020) | 54,244 | 1,431 | 1.6 | 1,431<sup>*a*</sup> | 1.00 | 1.6 | A | n | 0.026<sup>*e*</sup> | = |
| Crustaceans | Wolfe et al. (2019)<sup>*f*</sup> | 67,735 | 95 | 0.10 | 95 | 1.00 | 0.10 | Dt | — | 0.0014<sup>*e*</sup> | = |
| *Other animals* |  |  |  |  |  |  |  |  |  |  |  |
| Cephalopods | Basava et al. (2024)<sup>*c*</sup> | 868 | 81 | 9.2 | 81 | 1.00 | 9.2 | A | s | 0.093 | = |
| Bryozoa | Orr et al. (2022) | 6,613 | 565 | 5.1 | 565 | 1.00 | 5.1 | Dn | — | 0.085<sup>*e*</sup> | = |
| Cnidaria | Vaga et al. (2025)<sup>*f*</sup> | 11,597 | 289 | 2.2 | 289 | 1.00 | 2.2 | A | i | 0.025<sup>*e*</sup> | = |
| Sponges | Lavrov et al. (2023)<sup>*f*</sup> | 9,864 | 119 | 0.87 | 119 | 1.00 | 0.87 | A | i | 0.012<sup>*e*</sup> | = |
| Echinoderms | Mongiardino Koch et al. (2022)<sup>*f*</sup> | 7,749 | 65 | 0.80 | 65 | 1.00 | 0.80 | A | i | 0.0084 | = |
| Bivalves | Pfeiffer et al. (2019)<sup>*f*</sup> | 10,043 | 73 | 0.44 | 73 | 1.00 | 0.44 | N | — | 0.0073<sup>*e*</sup> | = |
| Nematodes | Qing et al. (2025)<sup>*f*</sup> | 28,415 | 156 | 0.36 | 156 | 1.00 | 0.36 | Du | — | 0.0055<sup>*e*</sup> | = |
| Gastropods | Zapata et al. (2014)<sup>*f*</sup> | 76,288 | 40 | 0.05 | 40 | 1.00 | 0.05 | A | i | 0.00052 | = |
| *Microbes & protists* |  |  |  |  |  |  |  |  |  |  |  |
| Fungi | Li et al. (2021)<sup>*f*</sup> | 156,891 | 1,620 | 0.89 | 1,620 | 1.00 | 0.89 | S | — | 0.010<sup>*e*</sup> | = |
| Diatoms | Nakov et al. (2018) | 16,427 | 1,113 | 0.24 | 1,113 | 1.00 | 0.24 | A | s | 0.068<sup>*e*</sup> | = |

## Materials and methods

### Groups and reference trees

I divided described eukaryotic species into 60 groups: clades of the rank at which species-level phylogenies are usually published (for example Birds, Seed plants, Nematodes), and residual sets (for example Other arachnids) that hold the remaining species, so that every described eukaryote belongs to one group. Bacteria and Archaea form 2 further groups, counted in species clusters of the Genome Taxonomy Database (Parks et al., 2026) and kept outside every eukaryotic number. The four levels of coverage are counted on one tree per group, its reference tree (below), because only those trees have their sequenced tips listed; they therefore depend on the grouping, since splitting a group can let a second tree count and merging two groups counts only the larger of their trees. The number of species named in any tree found does not depend on it: a species counts once if any tree names it. Both, and the size of the dependence, are given in Results (Coverage at four levels). Each group has the number of described species given by the source named in Supplement S1: a taxonomic authority or a published compilation, or the Catalogue of Life release COL26.9 where the count is attributed to the Catalogue. Some counts from compilations are rounded figures. The totals sum to 2,059,063 species, the denominator of every percentage in this paper.

For each group I took as its reference tree the largest published species-level tree I found, counting tips placed by taxonomy as well as by molecular data, whether the tree covers the whole group or one of its subclades. I searched Google Scholar, Dryad, Zenodo, Figshare, GitHub, TreeBASE, bioRxiv and clade portals (VertLife, FishTreeOfLife, insectphylo) up to May 2026, without a fixed query protocol. The rule is applied to the trees this search found; larger trees of several groups that it missed are not counted (Table 1 and Limitations). The species-level trees found, 263 with the reference trees of Bacteria and Archaea, are the catalogue of this paper (Supplement S1; Data availability); a group has no reference tree when the search found no downloadable species-level tree of it. Where a source distributes several trees, I used the one it designates as representative; otherwise the study’s chronogram where it covers every tip of the study’s other tree (Bryophytes, Gastropods and Turtles), the first tree of the fish all-taxon set (Rabosky et al., 2018), the diatom tree with more species carrying usable branch lengths (Nakov et al., 2018), and the maximum-clade-credibility tree where a source archives only a set of dated trees, computed with phangorn (Schliep, 2011). For trees that are updated continuously I used AvesData 1.3 (Birds), FTOL v1.8.0 (Ferns) and insectphylo Insecta_v2 (Insects). The bird release holds one synthesis aligned to successive versions of the Clements taxonomy up to Clements 2023, and for Birds the largest-tree rule is not the rule applied. The search took the Clements 2021 version; the Clements 2023 version has more tips (11,017) but fewer placed by an input phylogeny (9,189, against the sequenced tips of the bird reference tree in Table 1). For a synthesis aligned to several versions of one taxonomy, the version used is the one with the most tips placed by an input phylogeny, the tips that molecular coverage counts; the source’s crosswalk to the Open Tree taxonomy, through which the synthesis records the input trees of each tip, covers every tip of the Clements 2021 version and not every tip of the later versions (McTavish et al., 2025). Table 1 marks the Clements 2023 version; Supplement S1 states on which tree its phylogeny-placed tips are counted. I removed tips outside the group (outgroups and misplaced taxa) and kept one tip per species; tips labelled cf., aff., sp. or as hybrids are kept as unresolved labels (Supplement S2).

### Continuous data collection

Species-level trees continue to be published after the search, so the catalogue is maintained by a procedure that runs on a fixed schedule and changes no count by itself (its code is part of the data release; Fig. 1).

**Figure 1.**
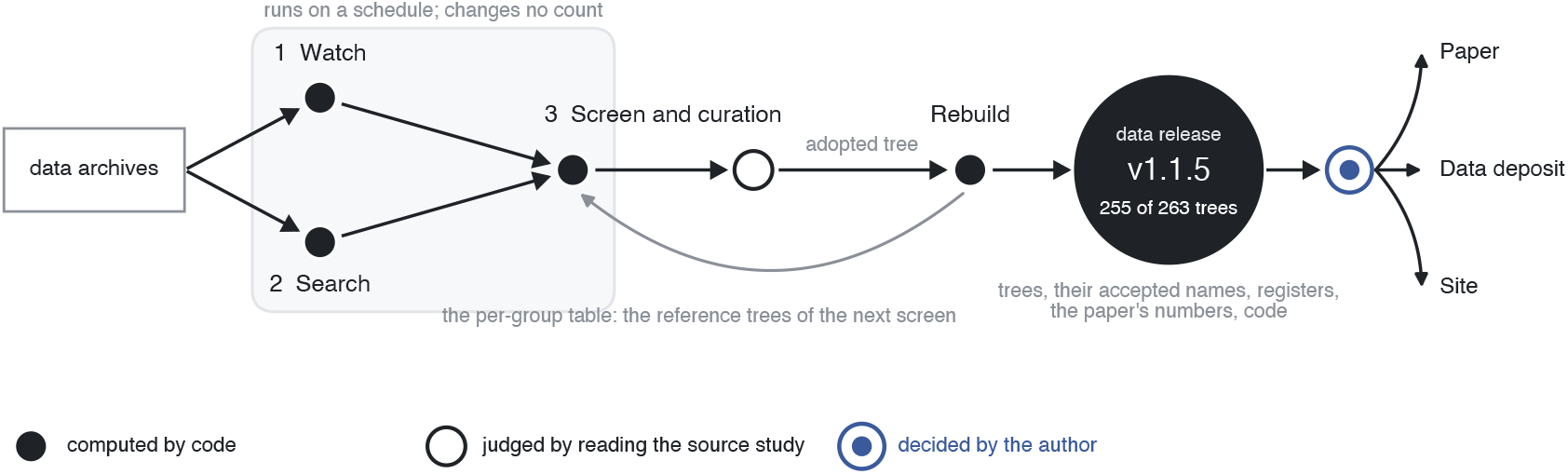
The procedure that keeps the catalogue current. (Materials and methods, Continuous data collection). Filled: computed by code; open: judged by reading the source study; blue: decided by the author.

#### Watch

The records deposited in Zenodo, Figshare and Dryad since the previous run whose metadata names a phylogeny, a chronogram or divergence times are collected, and every tree file they hold is read, inside archives too. Each tree is measured with the rules of this paper: tip labels are reduced to one key per species (Supplement S2), each genus is classified in the Catalogue of Life release COL26.9, and each species is assigned to the group whose definition contains it. The number of records each repository reports is compared with the number collected, and a run that misses records is marked incomplete.

#### Search

Each run also searches the whole history of the three repositories, group by group, with the group’s Catalogue of Life taxa and the same phylogeny terms, so that trees published before the watch began are found as well. A record is processed once: a ledger of every record seen keeps a later run from reading it again.

#### Screen and curation

A tree becomes a candidate when it is species-level, most of its species fall in one group, and it is at least nine tenths the size of the corresponding reference tree, or the group has none and the tree holds at least twenty of its species. Each group has two reference trees: the largest species-level tree dated in absolute time, and the largest species-level tree of any kind, which is the same tree where the largest tree is dated. A candidate is compared with each separately, so a larger undated tree never replaces a dated one. Curation then reads the source study for what cannot be measured: which tips were placed with sequence data, how the tree was dated, what date uncertainty its archive carries, and whether another file of the deposit is the tree to use; each verdict is recorded with its rule and a quotation from the source. An adopted tree enters a rebuild, which regenerates every number of the paper and the per-group table from which the next screen takes its reference trees, and produces a new data release; I approve each release before it is published as this article, the data deposit and the catalogue’s site. The counts of this version use the single reference tree defined above, on data release v1.1.5.

### Names

I resolved every tip label to an accepted species of the Catalogue of Life base release COL26.9 (ChecklistBank dataset 316321) (Bánki et al., 2026), counting synonyms of one accepted species once. The base release is assembled from the checklists the Catalogue curates; its extended release adds names from other sources, mainly GBIF, for groups those checklists cover incompletely. A label that carries no binomial, such as a lineage code, resolves only through a register row that gives the binomial the source’s own table of codes assigns to it (Supplement S3). Of the 481,229 tip labels, 11.5% resolve to no accepted species (listed per group in Supplement S3). Unresolved labels and tips merged into one accepted species together amount to 85,712 tips, 69,837 of them in the seed-plant tree, so a tree names fewer species than it has tips. As a sensitivity, I resolved the labels the Catalogue leaves unmatched against the GBIF backbone taxonomy (GBIF Secretariat, 2023), queried on 7–8 October 2026.

### Four levels of coverage

Each level is a number of described species divided by the group’s described species; eukaryotic totals sum numerators and denominators over groups, and each level is contained in the one before. The described species are also set against estimates of the total number of species (Results). A group’s estimate is a figure a published source states for the total number of its species, recorded as low, central and high values; where no source states one, the estimate is the described count, and where a source states only a range, the central value is the geometric midpoint of the range. Each group’s values and sources are in the data release.

*Named coverage* is the number of distinct accepted species among the tips of the reference tree.

*Molecular coverage* is the number of distinct accepted species among the tips that are in the molecular analysis that built the tree, here called sequenced tips. In 21 groups I listed these tips by intersecting the reference tree with the source’s own tree of sequenced species, alignment or list of sequenced taxa (Supplement S4). In the other 3 groups (Crocodilians, Spiders, Turtles) the source states that every tip has sequence data. A tip is sequenced when it is in the sequence analysis at all, whatever the number of loci behind it; the other tips of a tree are its taxonomically placed tips. For the bird tree, an automated synthesis that ran no sequence analysis of its own (McTavish et al., 2025), the sequenced tips are those of the source’s own tree of tips placed by an input phylogeny, the same rule as for the other groups’ trees of sequenced species; the tree of Jetz et al. (2012) among the inputs is their tree of species with genetic data. The source’s tree also holds tips whose only input is one of the taxonomy trees of the synthesis, or whose inputs could not be matched; leaving them out gives the lower end of the bird fraction, a sensitivity (Supplement S4). The molecular fraction of a tree is the share of its tips that are sequenced; it differs from the molecular sampling fraction, which divides sequenced tips by described species. The two sampling fractions of Table 1, tips and sequenced tips over described species, count tips and so assume one described species per tip; Table 1 marks the groups in which tips exceed named species by more than 10% (Names). *Dated coverage* is molecular coverage in reference trees that are chronograms in absolute time: branch lengths in millions of years and all tips at the present, the second measured on each file as a coefficient of variation of root-to-tip distances of at most 0.05 (Supplement S5). Ultrametric trees in relative time do not count. For each tree that is not a chronogram, I recorded from the source study whether it dated the tree and archived the result.

The source archive of a tree is the set of files its authors archived with the study. *Date uncertainty available* is dated coverage in trees whose source archive carries variation in node dates that a user can propagate: a tree set whose node ages vary among the sequenced species (posterior sets, bootstrap datings, sets with dates drawn from published studies or with root ages drawn from a calibration prior), or intervals annotated on the nodes of a summary tree. It counts archived variation in dates, whatever its source and its relation to the calibration priors. A tree set that adds taxonomically placed species to one dated tree does not count: pruned to its sequenced tips, each of its trees has the dates of the source’s one dated tree (the fish set of Rabosky et al. (2018); Supplement S6). Whether an archived interval was narrowed by the sequence data beyond what the calibrations allow is not a level: answering it requires a run of the source’s analysis that samples from the prior, which no archive holds; I made one for Cephalopods from the source’s input file, and for no other group. Supplement S6 describes, for each archived crown interval (the node joining the living members of the clade the source study sampled), its comparison with the range its calibrations allow, or that it has no range to compare with. Where a source archives the set of dated trees that its summary tree summarises, the interval is read from the set. The sets of Amphibians, Mammals and Squamates also place species without sequence data. Each source dated a backbone of one representative per subclade in MrBayes under fossil or secondary calibrations, and each tree of its set carries one tree of that backbone posterior, with the subclade trees rescaled to it and the species without sequence data placed within the subclades (Jetz and Pyron, 2018; Upham et al., 2019; Tonini et al., 2016). The crown is a backbone node, so its spread over the set is the posterior of the backbone dating.

### Sensitivity analyses

I recomputed the four levels under these alternatives:

- denominators: restricted to groups with a tree; a Catalogue of Life count for every group (Bánki et al., 2026);
- names and trees: the named species of all trees of the catalogue added, with an upper bound for molecular coverage that counts every added species as sequenced; names resolved with GBIF added; the bird tree removed;
- molecular coverage computed as named species times the molecular fraction of tips;
- date uncertainty counted only where the archive holds dated trees of a posterior or bootstrap dating of sequence data whose crown age is neither fixed in every tree nor at its median on a hard bound (Table 2).

**Table 2.**
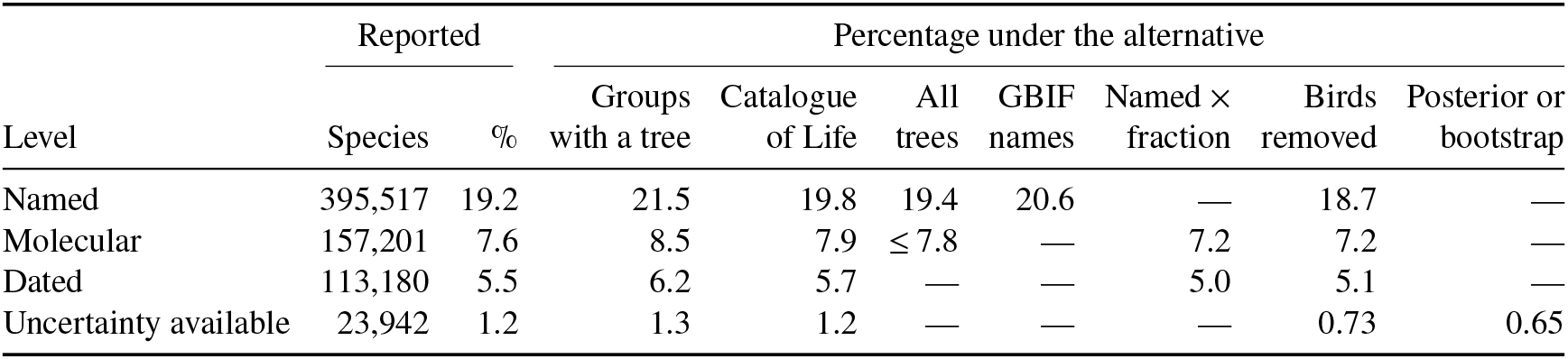
Coverage at the four levels and its sensitivity. Species and percentage of the 2,059,063 described eukaryotic species, then the percentage under each alternative: the denominator restricted to the 24 groups with a tree (1,839,402 species); a Catalogue of Life count for every group (1,993,667 species); named species of the other 237 eukaryotic trees of the catalogue added, with an upper bound for molecular coverage that treats every added species as sequenced; tips that the Catalogue leaves unmatched resolved with the GBIF backbone; molecular coverage computed as named species times the molecular fraction of tips; the automated-synthesis bird tree removed; date uncertainty counted only where the archive holds posterior or bootstrap datings of sequence data whose crown age is neither fixed nor on a bound (Amphibians, Cephalopods, Mammals, Squamates and Turtles). A dash marks an alternative that does not apply to that level.

Further sensitivities are in Supplement S7.

### Comparison with the Open Tree of Life

From the Open Tree of Life synthesis release v16.1 I took the extant eukaryotic species placed by at least one input phylogeny (Hinchliff et al., 2015). I compared them with the sequenced species of the reference trees by exact genus and epithet, and again after resolving the Open Tree names to Catalogue of Life accepted species by the rule used for the tips, so that synonyms are matched (Supplement S8). The comparison tells a user whether the Open Tree’s phylogeny-placed species can stand in for the sequenced tips of a reference tree.

### Worked example

Under a constant-rate birth–death model with uniform sampling, the likelihood of a tree depends on the speciation rate *λ* and the sampling fraction *ρ* (sampled over described species) only through their product and through net diversification *r* = *λ* − *μ* (Stadler, 2009). Changing *ρ* on a fixed tree therefore leaves *r* unchanged. It multiplies *λ* by the ratio of the old to the new fraction and changes extinction *μ* and the extinction fraction *ε* = *μ*/*λ* with it, unless that would require *μ* < 0; then the maximum-likelihood estimate lies on the boundary *μ* = 0 and the likelihood falls (Cusimano and Renner, 2010; Höhna et al., 2011). Extinction is estimated with little precision from trees of living species (Nee et al., 1994; Rabosky, 2010), and if rates vary through time, constant-rate estimates of *μ* and *ε* have no unique interpretation (Louca and Pennell, 2020). The direction of the change is known before any fit; the example measures its size and finds which trees reach *μ* = 0.

I fitted the model to every reference tree that is dated in absolute time and has fewer listed sequenced tips than tips (7 groups: Amphibians, Birds, Fish, Mammals, Seed plants, Sharks and Squamates). Each tree is fitted with the branch lengths its source archived; after outgroups are removed its root is the crown. Where the source summarises the crown age over a tree set, that age differs from the root height of the summary tree by factors from 0.98 to 1.04 (crown age over root height); the trees are not rescaled to it. Multiplying every branch length by a constant divides *r*, *λ* and *μ* by it and leaves *ε* and every likelihood-ratio statistic unchanged, so the rates are on the time axis of the archived tree. The trees of Amphibians, Seed plants and Squamates have nodes with more than two children, left by a consensus summary or by species attached by taxonomy to an unresolved node; the likelihood reads a node with *k* children as *k* − 1 speciation events at its age, so these events are not branching times estimated from sequence data, in the pruned tree as in the full one. I report the share of events at such nodes. Each tree was fitted three times: the full tree with *ρ* equal to its tips over described species; the tree pruned to its sequenced tips with *ρ* equal to sequenced tips over described species, the fraction that matches it; and the same pruned tree with the full tree’s fraction, as a user would if they pruned a downloaded tree and kept the fraction quoted for it. I report 95% profile-likelihood intervals, conditional on the one tree fitted; where the estimate of *ε* is zero, the interval runs from zero to its upper end. Write *ρ_m_* for the matching fraction and *ρ_f_* for the full tree’s fraction. The fit with *ρ_f_* is the matching model restricted to *ε* ≥ 1 − *ρ_m_*/*ρ_f_*, the values for which extinction under *ρ_f_* is not negative. I tested this restriction with the likelihood-ratio statistic, twice the difference of the two maximised log-likelihoods. The null hypothesis is joint: constant rates, uniform random sampling of the sequenced tips with fraction *ρ_f_*, and *μ* ≥ 0. Sequenced tips are not a uniform random sample of a group’s species, and clustered or diversified sampling lowers estimated extinction (Cusimano and Renner, 2010; Höhna et al., 2011), which can produce a rejection whatever the fraction; a rejection therefore does not attribute the misfit to the fraction alone. The full tree’s fraction does not match the pruned tree, so the joint null is false by construction; the test asks whether a tree can show it. Where the matching estimate of *ε* already satisfies the restriction, the two fits have the same likelihood on that tree and the statistic is zero; a zero statistic carries no information about whether the fraction is correct. Whether the restriction binds is a property of the tree fitted and can differ between trees of one group. At the edge of the restriction the statistic follows an equal mixture of a point mass at zero and a chi-squared distribution with one degree of freedom (Self and Liang, 1987); its critical value at the 5% level is 2.71, and *p*-values are computed under that mixture. A test is reported only where its model assumptions hold for the tree: the tree is binary, and its node ages are estimated by a dating analysis. The speciation events that the likelihood reads at one age, at a node with more than two children, have probability zero under the model, and on the consensus trees the statistic moves when the polytomies are resolved at random (Supplement S9), so on a summary tree with polytomies (Amphibians, Seed plants and Squamates) the statistic is reported without a test or *p*-value. The bird summary tree assigns published ages to the nodes the source studies share and interpolates the others, and its tree of the tips placed by input phylogenies, before taxon addition, is not binary (McTavish et al., 2025); its statistic (Birds) is reported without a test, with the share of its node ages that are not published dates and the number of its nodes no input phylogeny resolves (Supplement S9). Both conditions are read from the trees and the source files before any fit, and no fitted quantity enters them. The family of tests is the 3 fitted groups whose pruned summary tree meets both conditions (Fish, Mammals and Sharks); each test is made at the Bonferroni level 0.0167 (5% over 3 tests), with critical value 4.53. So that the outcome of the other choice can be read, the bird statistic is also compared with the critical value of a family of 4 tests that includes it (level 0.0125, critical value 5.02). Where a source archives the dated trees that its summary tree summarises, each tree binary and dated in the source’s analysis, I repeated the three uniform-sampling fits on an evenly spaced sample of them, pruned to the same sequenced tips (Amphibians, Birds, Mammals, Sharks and Squamates; Supplement S9), so that the summary-tree estimate is one point of a distribution. The trees of a set are draws from one posterior, one procedure of date assignment or one prior on the root age, not independent replicates, so I describe the distribution of the statistic over the set, by its median, its range and the number of trees in which it exceeds each critical value, and form no test from it: a number of trees above a critical value has no error rate, and a median of per-tree statistics has no reference distribution under the null. For the groups whose source archives a posterior set (Amphibians, Mammals and Squamates), that distribution is the primary result and the summary-tree test is secondary. For Sharks, both archived sets vary node ages through the root age (Methods, ‘Temporal Calibration’ and ‘Taxon-complete trees’; Stein et al., 2018): 500 root ages were drawn from the lognormal calibration density of the root; the maximum-likelihood tree of the 610 sequenced species, its topology fixed, was dated in treePL under each of these root ages; the other 582 species were added to each dated tree within taxonomic constraints and resolved with Polytomy Resolver, which samples their placements in BEAST from the prior only; and 20 draws for each of the 500 root ages form the 10,000 trees of the set. Over these sets the statistic describes the sensitivity of the outcome to the root-age draw, not a distribution over a dating posterior. I report beside the summary tree the distributions over the source’s dated trees of its sequenced species and over its taxon-complete trees pruned to the same species. To find why these two differ, I compared each taxon-complete tree fitted, pruned to the sequenced species, with the source’s trees of those species: clades matched by their tips, the node ages of matched clades, and the number of nodes younger than 5 Myr (Supplement S9). For Fish and Seed plants the summary tree carries the result. For Mammals and Sharks, whose sources also archive dated trees of their sequenced species alone, I fitted a sample of those trees with the matching fraction. As a check of the consensus summaries only, every uniform-sampling fit was also repeated with the polytomies resolved at random. Because uniform sampling is an assumption, I refitted each pruned tree with tip-specific sampling fractions, each tip carrying the fraction of its family (the family’s sequenced species over its accepted species in the Catalogue of Life). The likelihood carries the probability that a lineage leaves no sampled descendant along each branch from that branch’s tip fractions, as in the treatment of incomplete sampling of FitzJohn et al. (2009) and the clade-specific fractions of BAMM (Rabosky, 2014); unsampled species are attached to no clade, so families need not be monophyletic, and the species of families with no sequenced tip enter no fraction. Where the two children of a node carry different values, the node takes the value of one child, so this likelihood is an approximation whose value depends on the order in which children are stored (Moore et al., 2016); I report the fit with that order reversed. The family and uniform fits make different sampling assumptions and are not nested, so their likelihoods are not compared; I report the estimates side by side. Every fit, the family fractions and a check of the likelihood against phytools are in Supplement S9.

### Software and data

Counts are computed in Python 3.14; the birth–death fits use R 4.5.1 with ape 5.8.1 (Paradis and Schliep, 2019), and the maximum-clade-credibility trees use phangorn 2.12.1. One script regenerates every derived file and table, and the data of every figure, from the archived trees and registers (Supplement S10).

## Results

### The reference trees

Of the 60 eukaryotic groups, 24 have a species-level reference tree and 36 have none (Fig. 2). The groups without a tree hold 10.7% of described eukaryotic species; the largest of them are Acari, Flatworms, Annelids and Myriapods. Of the 24 reference trees, 18 are chronograms in absolute time (Table 1). Of the others, the insect tree is ultrametric in relative time; the fungal tree is in substitutions and its study archived a dated tree in relative time only; the bryozoan study did not archive its chronogram; the crustacean study archived one that lacks some tips; the nematode study archived one that could not be read; and the bivalve study did not date its tree.

**Figure 2.**
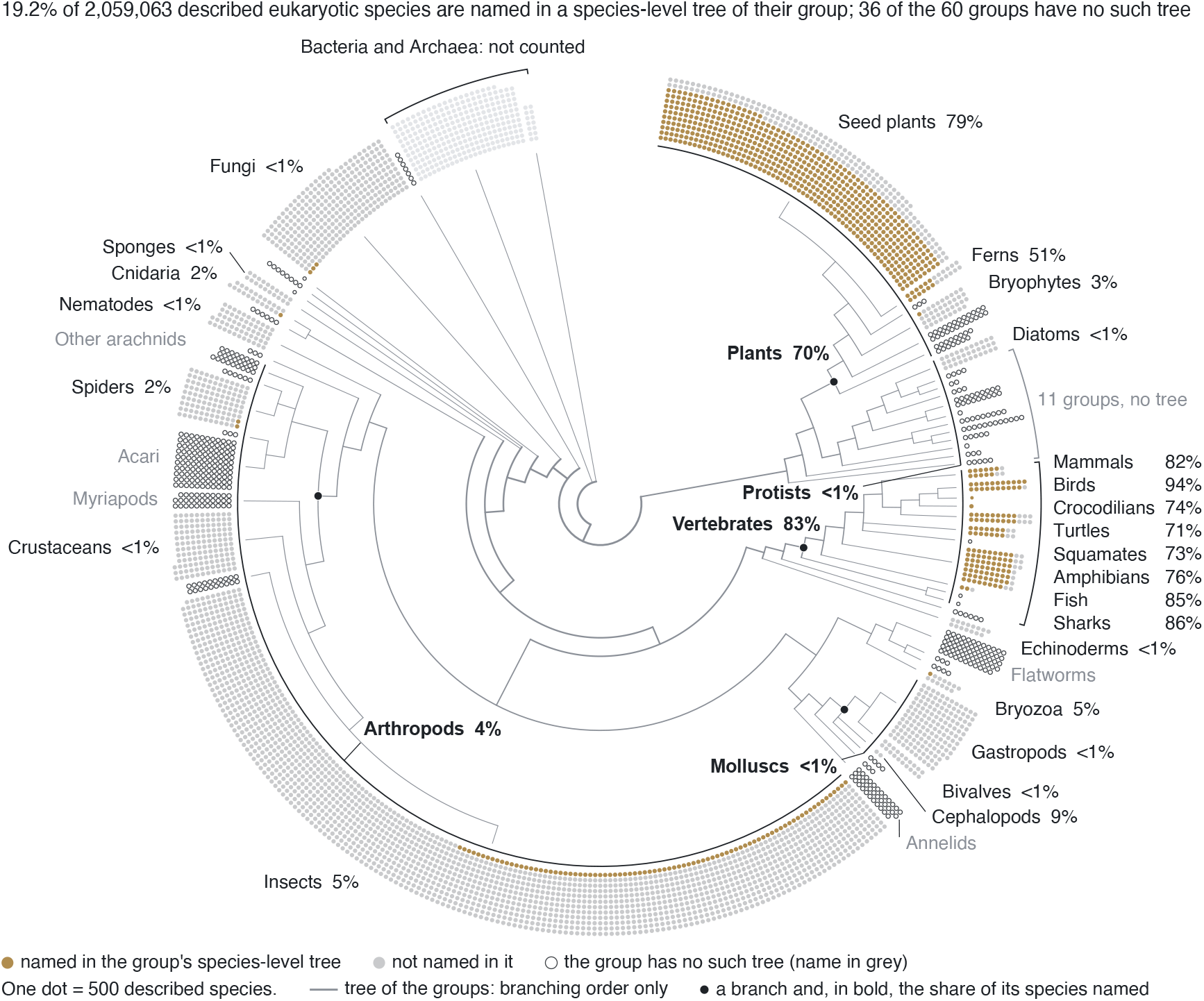
The groups around the tree of life, with the share of their described species named in a reference tree, as drawn by the catalogue’s site. (https://franciscorichter.github.io/phylo-species-atlas/). Each dot stands for five hundred described species; each arc is one group, as wide as its described species (the 60 eukaryotic groups, and Bacteria and Archaea), and the arcs follow the branching order of the groups in the Open Tree of Life synthesis, drawn inside the ring as a tree that shows that order only. Brass dots: species named in the group’s reference tree, the species-level tree of the key; grey dots: species of a group with a reference tree that the tree does not name; open circles: the 36 groups with no species-level tree, their names in grey. A group smaller than one dot is drawn as one small mark, and a named count below half a dot draws no brass dot. The percentage after a group’s name is its named species over its described species. Arcs along the ring mark five sets of groups, named in bold with the share of their species named: four are branches of the tree, each marked by a dot, and Protists is a run of groups that is not one branch. The vertebrate groups are listed beside their bracket in ring order. The smaller groups are not labelled; clockwise from the top they are Lycophytes, Green algae, Rhodophyta, Brown algae, Chrysophyta, Oomycetes, Foraminifera, Cercozoa, Ciliates, Apicomplexa, Dinoflagellates, Haptophytes, Euglenozoa, Amoebozoa, Other vertebrates, Lampreys, Hagfish, Tunicates, Acanthocephala, Rotifera, Other Mollusca, Nemerteans, Other hexapods, Pycnogonida, Scorpiones, Tardigrada, Myxozoa, Ctenophores, Other phyla, Choanoflagellates and Other protists; the grey bracket spans eleven of them, Brown algae to Amoebozoa. The catch-all groups (Other vertebrates, Other Mollusca, Other arachnids, Other phyla, Other protists) are not clades; each sits at the common ancestor of its members. Bacteria and Archaea (the palest dots, under one bracket; GTDB species clusters) are drawn apart and are outside every count.

### Coverage at four levels

The 24 reference trees and the 237 other trees of the catalogue name 399,548 described eukaryotic species (19.4%); the reference trees alone name 395,517 (19.2%, named coverage; Fig. 3a). Of the species the reference trees name, 157,201 (7.6%) are placed with molecular data and 113,180 (5.5%) are in a tree dated in absolute time. Date uncertainty is available for 23,942 species (1.2%), 38% of them in the tree of Birds; where the archive holds posterior or bootstrap datings of sequence data with a free crown age (Amphibians, Cephalopods, Mammals, Squamates and Turtles), it is available for 13,370 species (0.65%; Table 2). Molecular coverage is 39.7% of named coverage and dated coverage 72.0% of molecular coverage; date uncertainty is available for 21.2% of dated coverage, and posterior or bootstrap datings

**Figure 3.**
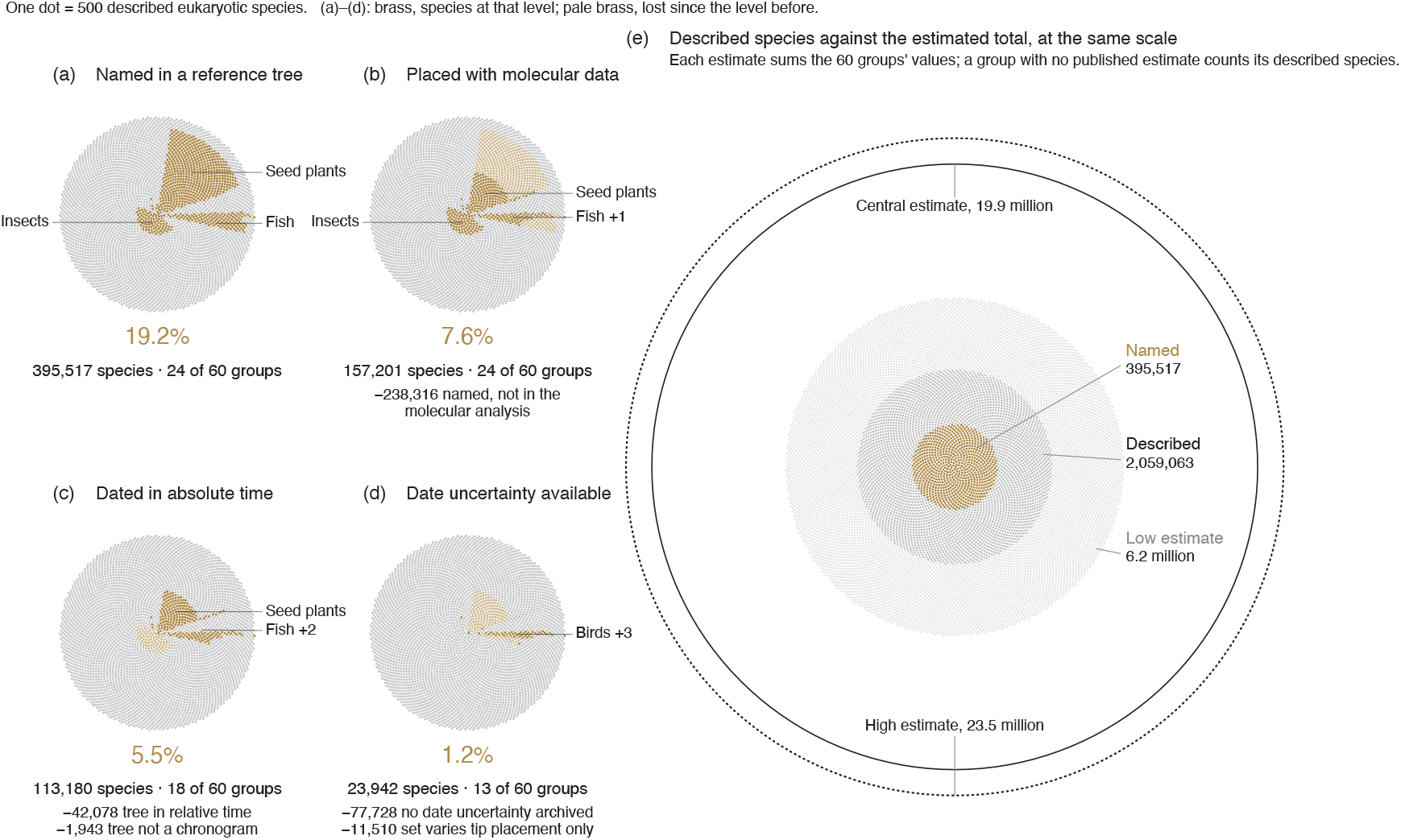
Described eukaryotic species at four levels of coverage, and against the estimated number of species, as drawn by the catalogue’s site. (a–d) Each disc holds the 2,059,063 described eukaryotic species, one dot per five hundred species, in the 60 groups in tree-of-life order clockwise from the top; within each group the lit dots are drawn nearest the centre. Brass dots are the species (a) named in a reference tree, (b) placed with molecular data, (c) in a tree dated in absolute time and (d) in a dated tree whose archive carries date uncertainty; pale brass dots were lit at the previous level and lost at this one. Percentages are shares of described species. Below each disc: the percentage, the count, the groups that contribute, and the species lost from the previous level by reason. The groups contributing most are named; “+N” counts further groups in the same sector: Birds in (b); Ferns and Birds in (c); Mammals, Squamates and Amphibians in (d). (e) The same dots at the same density: the species named in a reference tree (brass) inside the described species (grey), faint dots out to the sum of the groups’ low estimates, a thin circle at the sum of their central estimates and a dotted circle at the sum of their high estimates (Results); a group with no published estimate counts its described species. Bacteria and Archaea are not shown. of sequence data with a free crown age for 11.8%. With the classes, orders or families of the Catalogue of Life as the groups, the count with one tree per group is within 1.1% of the count over every tree, for named and dated species and for species with date uncertainty available; with phyla as the groups it falls to 88.8% of it for named species and 33.4% for date uncertainty, because one tree then stands for a whole phylum (Supplement S7). The groups’ estimates of their total number of species sum to 6.2 million (low), 19.9 million (central) and 23.5 million (high; Fig. 3e); Nematodes alone holds 50.2% of the central sum. Across the alternatives named coverage ranges from 18.7% with the bird tree removed to 21.5% over groups with a tree, and molecular coverage from 7.2% to 8.5% (Table 2). The other trees of the catalogue add 4,031 named species. Counting every tip as a species, an upper bound that ignores unresolved and merged labels, gives 23.4%.

### Coverage by group

Coverage is concentrated in a few groups (Fig. 2). Vertebrate trees include 83.0% of described vertebrates and place 44.4% with molecular data, and plant trees include 70.3% and place 19.0%. Arthropod trees include 3.7%, and those of other animals and of protists and fungi 0.46% and 0.69%. The seed-plant tree (Smith and Brown, 2018) alone holds 70.6% of the named species, 45.0% of the sequenced species and 62.5% of the dated species. Without that tree and the described seed plants, eukaryotic named coverage is 6.8%, molecular coverage 5.1% and dated coverage 2.5%. In the diatom tree many labels are strain or sample names without a binomial, and most of the binomials are absent from the Catalogue’s base release, which lists far fewer diatom species than the group’s authority (Supplement S3 and S7), so for diatoms the name authority, not the tree, sets named coverage.

### Named against molecular coverage

Among the trees that include the largest share of their groups are the two with the lowest molecular fractions (sequenced tips over tips), Seed plants (0.22) and Fish (0.37), in which most tips were attached by taxonomy (Fig. 4). Named coverage of Seed plants is 78.8% and molecular coverage 20.0%; for Fish the two are 85.4% and 31.7%. Taking the largest tree favours trees completed by taxonomy, so this pattern follows partly from the selection rule. The bird tree, the only automated synthesis, has a molecular fraction of 0.85. For all 7 trees that contain taxonomically placed tips, the source also distributes a tree of its sequenced species (Table 1; Supplement S4). Measured on the distributed files, 4 of these are dated (Fish, Mammals, Seed plants and Sharks), those of Amphibians and Squamates are maximum-likelihood trees with branch lengths in substitutions, and that of Birds has no branch lengths.

**Figure 4.**
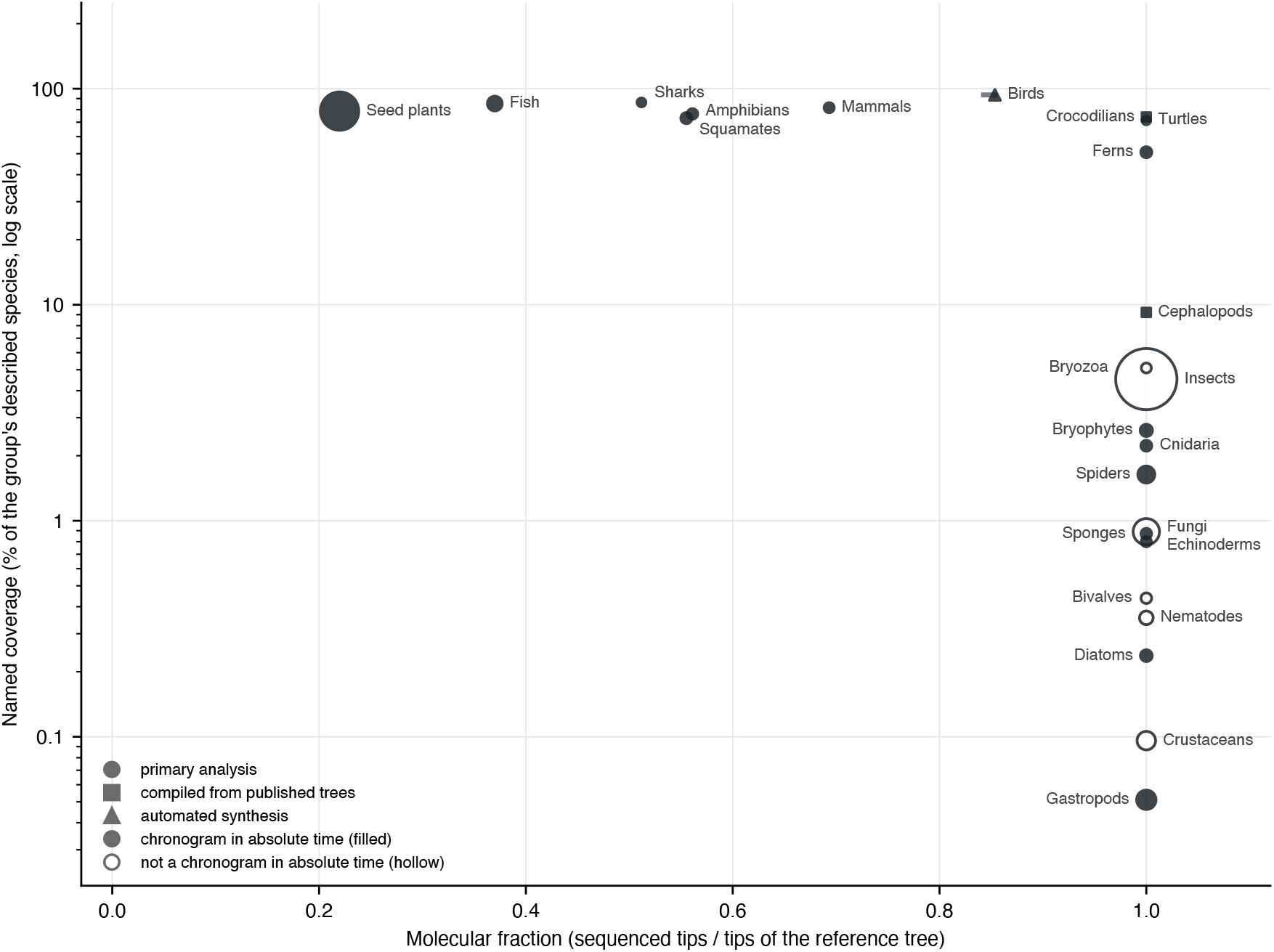
Named coverage against the molecular fraction of each reference tree. Each point is one of the 24 reference trees; the horizontal axis is the share of its tips that are in the molecular analysis behind it, and the vertical axis its named coverage on a logarithmic scale. Marker area: the group’s described species; filled: dated in absolute time. Shape: how the tree was assembled; a primary analysis is one study’s own inference from sequence data, whether the sequences were newly generated or mined from public databases; a tree compiled from published trees takes its topology from earlier studies (Crocodilians, and Cephalopods, whose study dated that topology with its own alignment); an automated synthesis combines published trees and taxonomy (Birds). A horizontal bar marks a range of molecular fractions.

The Open Tree of Life places 129,889 extant eukaryotic species by at least one input phylogeny (6.3% of described species). In the 21 groups with listed sequenced tips, the reference trees place 156,037 accepted species with molecular data, of which 74,000 (47.4%) are among the Open Tree’s phylogeny-placed species after both are resolved to accepted names. Matching by exact spelling finds 74,935 of the 174,256 names of the sequenced tips (43.0%) among the Open Tree’s phylogeny-placed names, so synonymy does not explain the difference; in the other direction, the same 74,935 names are 59.0% of the 126,904 distinct names of the 128,032 phylogeny-placed species of these groups (Supplement S8). 82,037 sequenced species of the reference trees are absent from the Open Tree’s phylogeny-placed set. Of these, 46.3% are in the seed-plant tree and 41.4% in the insect tree, both built from sequences mined from public databases (Smith and Brown, 2018; Chesters et al., 2023); whether their species are missing from the Open Tree’s input studies was not tested (per group in Supplement S8).

### Where dating and date uncertainty are lost

Of molecular coverage, 28.0% is in trees that are not dated in absolute time, and 95.6% of that is in the insect tree (42,078 species; Fig. 3c). Of dated coverage, 68.7% is in chronograms archived with no interval or tree set (Crocodilians, Ferns, Seed plants and Spiders; 77,728 species), and 10.2% (11,510 species) in a chronogram whose tree set varies only the placement of the taxonomically added tips (Fish), so date uncertainty is not available for them.

Of the 18 dated reference trees, 14 carry an interval of some kind and 4 carry none (Fig. 5). Date uncertainty is available in 13 of them: 9 archive tree sets (Amphibians, Birds, Bryophytes, Cephalopods, Diatoms, Mammals, Sharks, Squamates and Turtles) and 4 summary trees with node intervals (Cnidaria, Echinoderms, Gastropods and Sponges). What the sets vary differs: the set of Birds draws its dates from published studies (McTavish et al., 2025), and that of Sharks draws its root ages from the calibration prior (Materials and methods). Of the archived intervals, 7 lie within the range their calibrations allow and are not narrowed below half its width; Supplement S6 gives the comparison for each interval.

**Figure 5.**
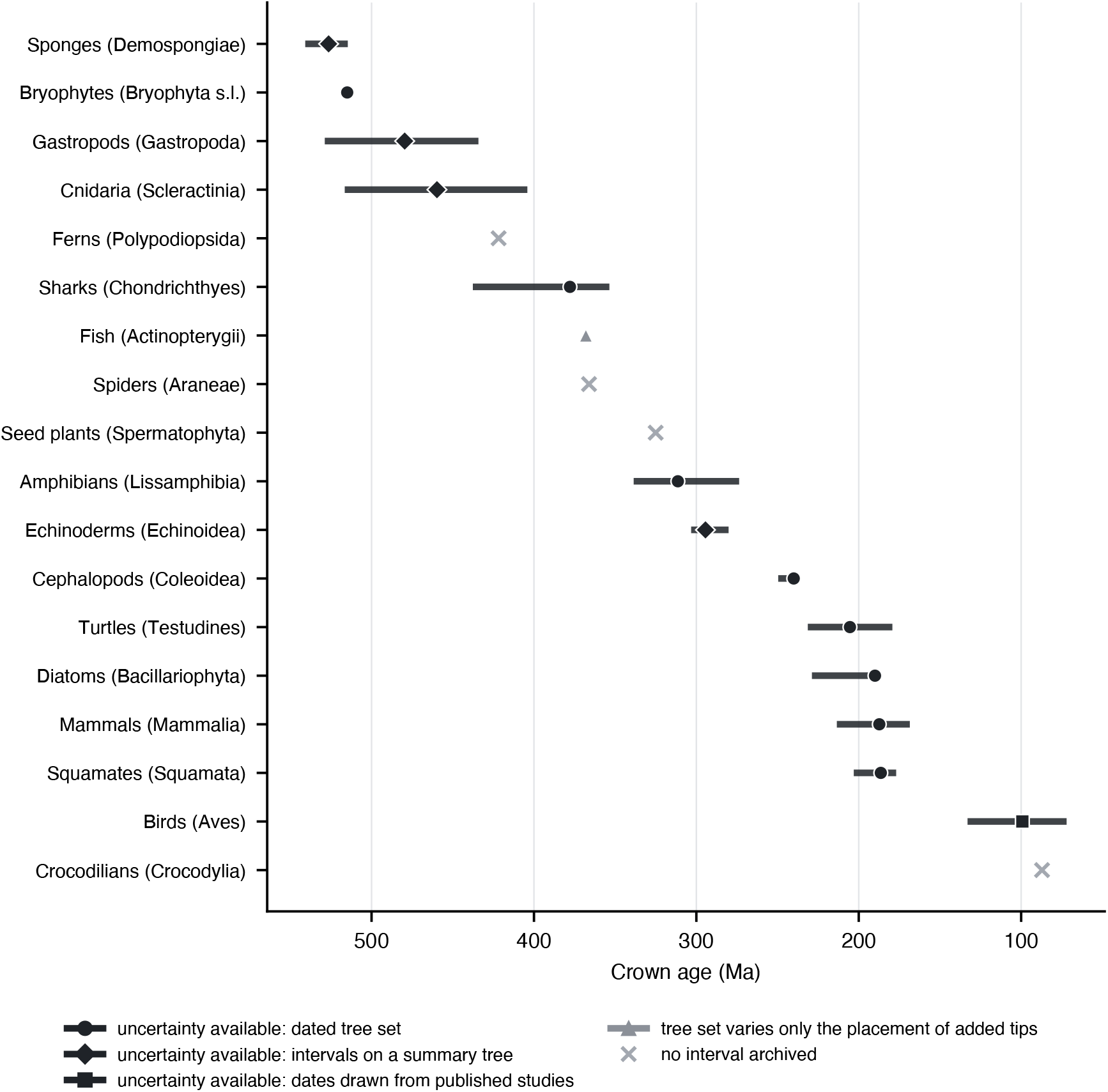
Crown ages and archived intervals of the 18 reference trees dated in absolute time. Points: crown age of the clade the source sampled (in parentheses), from the source’s own summary of the node; bars: the archived interval. Date uncertainty a user can propagate is archived as a dated tree set (circles), as intervals on the nodes of a summary tree (diamonds), or as a set whose dates are drawn from published studies (square). Triangle: the tree set varies only the placement of taxonomically added tips, and carries no date uncertainty for the sequenced species. Crosses: no interval is archived. Supplement S6 compares each crown interval with its calibrations.

### Worked example: sampling fraction and tree choice

On the tree pruned to its sequenced tips, replacing the matching sampling fraction by the full tree’s fraction lowers the speciation rate in all 7 groups, as the likelihood predicts (Table 3, Fig. 6). On the summary trees the restriction does not bind in 4 groups (Fish, Mammals, Seed plants, Sharks), so *λ* is rescaled and *r* unchanged (Materials and methods), and the extinction fraction falls, in Fish from 0.75 to 0.32 and in Sharks from 0.54 to 0.10. For Sharks the profile interval of the matching extinction fraction on the summary tree (0.40–0.65) includes the edge of the restriction, 1 − *ρ_m_*/*ρ_f_* = 0.49, and on the source’s dated trees of its 610 sequenced species the restriction binds in 84 of 100 trees, with a statistic of median 0.68 and range 0.0–16.8, above 2.71 in 3 of them and above the corrected critical value of 4.53 in 2. These trees share one topology and differ in node ages through the root age drawn for each, so the count measures the sensitivity of the outcome to the root-age draw, not its distribution over a dating posterior; under that draw the summary-tree outcome for Sharks does not hold on most of these trees. The source’s taxon-complete trees, pruned to the same 610 species, give a different distribution: the restriction binds in 7 of 100 trees, with a statistic of median 0.00 and range 0.0–4.4, above 2.71 in 1 of them and above 4.53 in 0, and the matching extinction fraction runs from 0.31 to 0.72, against 0.09 to 0.64 on the trees of the sequenced species. The two sets are not the same trees of the sequenced species. Every tree of the sequenced-species set has the source’s one maximum-likelihood topology; pruned to the sequenced species, none of the 100 taxon-complete trees compared has it, and each lacks from 111 to 142 of its 609 clades (median 129, 77 of them within one genus). Matched to the tree of the sequenced-species set whose clade ages are closest, a median of 307 clades have ages within 1%. The pruned taxon-complete trees have a median of 51 nodes younger than 5 Myr against 38, so their branching times are spread more evenly towards the present, which a constant-rate fit reads as higher extinction, above the edge of the restriction. The taxon addition with Polytomy Resolver, which samples the placement and age of each added species in BEAST from the prior, also changed the relationships and node ages of the sequenced species (Stein et al., 2018); the taxon-complete set is a set of complete trees, and the set of the sequenced species is the source’s dating of the sequence data under each root age drawn. The full tree’s fraction requires negative extinction in every squamate posterior tree fitted (100 of 100), in 89 of 100 amphibian posterior trees and in 0 of 100 mammal posterior trees; for Birds it does so on the summary tree, but in 2 of 100 trees with dates drawn at random and 0 of 100 with mean dates. Over the posterior tree sets (Amphibians, Mammals and Squamates; Table 3; Supplement S9), the statistic ranges from 2.0 to 90.7 (median 35.57) over the squamate posterior trees and from 0.0 to 20.5 (median 1.80) over the amphibian ones. Against the corrected critical value of 4.53, the groups whose statistic lies above it in every posterior tree: none; on both sides of it: Amphibians and Squamates; below it in every tree: Mammals. Over the amphibian posterior set it exceeds 4.53 in 27 of 100 trees and the uncorrected 2.71 in 42, so where one tree’s statistic lies depends on which posterior tree is fitted. The posterior sets are taxon-complete (Materials and methods), so the ages of the sequenced species in the pruned trees are not a dating of their sequence data alone. On the squamate posterior trees the matching extinction fraction ranges from 0.14 to 0.41, against 0.15 on the consensus tree. The summary trees of Amphibians and Squamates are consensus trees with 29.0% and 24.4% of the pruned tree’s branching events, and 48.5% and 44.7% of the full tree’s, at polytomies; on the full trees their estimates of *r* (0.034 and 0.057) and of *ε* lie outside the ranges over the posterior sets (Table 3; Supplement S9), and their profile intervals describe that one tree (Supplement S9). On the consensus trees the statistic is 8.16 for Amphibians and 83.74 for Squamates, and with their polytomies resolved at random it runs from 9.5 to 12.5 and from 58.8 to 63.0 (Supplement S9); these statistics carry no test. The restriction binds on none of the 3 summary trees tested (Fish, Mammals and Sharks), so their statistic is zero and no test of the family rejects. On the bird summary tree the restriction binds, with a statistic of 4.99, and no test is made, because its node ages are not estimated by a dating analysis: 3,836 of its 10,823 internal nodes (35.4%), and 2,251 of the 9,238 of the tree pruned to its sequenced tips (24.4%), carry no published age and are interpolated or dated at taxon addition (McTavish et al., 2025). Before taxon addition, the synthesis of the 9,239 tips an input phylogeny places has 9,120 internal nodes and 231 branching events at polytomies, so it is not binary; the binary pruned summary tree carries 118 nodes that the input phylogenies leave unresolved. In a family of 4 tests that includes the bird tree, at level 0.0125 with critical value 5.02, its statistic of 4.99 does not reject, so no test rejects under either family. On the bird sets, which vary the dates drawn from published studies and the placement of taxonomically added tips, the matching extinction fractions, from 0.10 to 0.27 and from 0.18 to 0.19, lie above the 0.07 of the summary tree; this comparison describes the bird trees and is not a criterion of the family. On the mammal source’s dated trees of its sequenced species, the matching extinction fraction ranges from 0.68 to 0.74, against 0.70 on the pruned reference tree, whose node ages were estimated together with the taxonomically placed species; with the full tree’s fraction the restriction binds in 0 of 100 of these trees.

**Figure 6.**
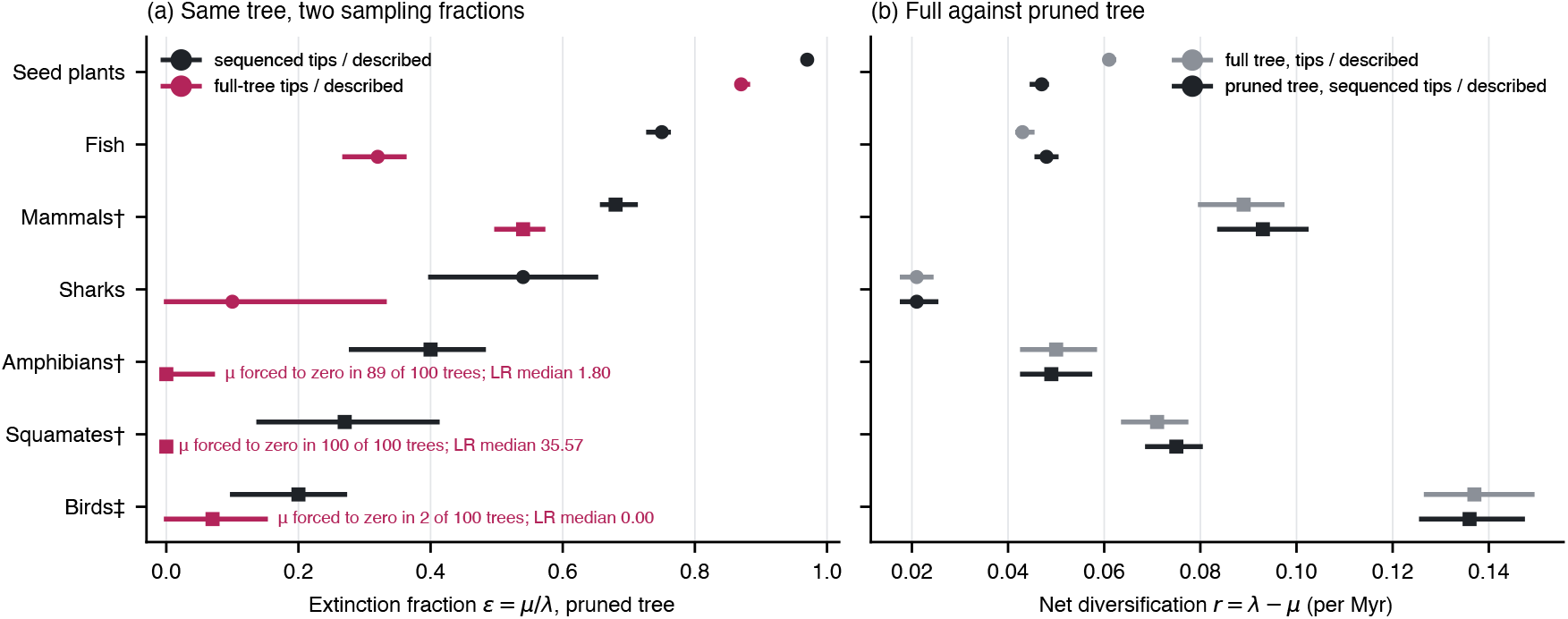
A constant-rate birth–death model fitted to the 7 dated reference trees that contain taxonomically placed tips. (a) Extinction fraction on the tree pruned to its sequenced tips, with the sampling fraction that matches that tree (sequenced tips over described species, ink) and with the full tree’s fraction (tips over described species, red). Where the full tree’s fraction forces extinction to zero over a set, the label gives the trees in which it does and the median likelihood-ratio statistic against the matching fit. (b) Net diversification rate on the full tree and on the pruned tree, each with its own sampling fraction. Squares, for Amphibians, Mammals and Squamates (†, the source’s posterior set) and Birds (‡, the set with dates drawn at random): median and range over the trees fitted of the set. Circles: summary-tree estimate with its 95% profile-likelihood interval, conditional on that one tree; an interval whose estimate is zero starts at zero. The summary-tree fits of the set groups are in Supplement S9.

**Table 3.**
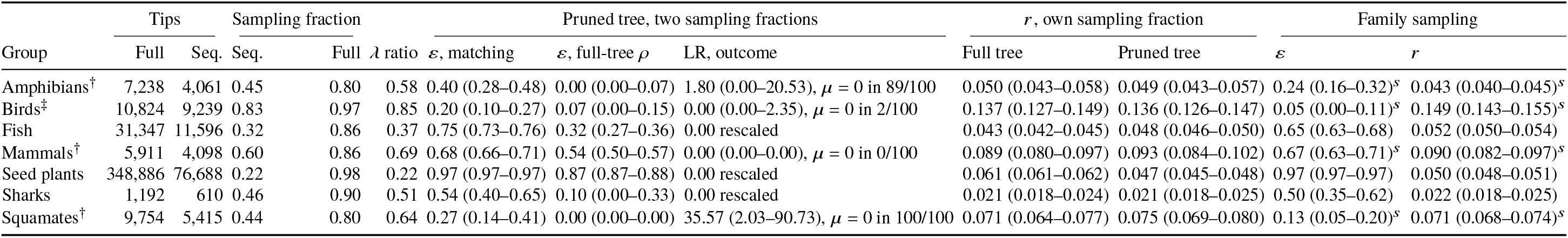
Constant-rate birth–death fits to the 7 dated reference trees that contain taxonomically placed tips. Sampling fractions: sequenced tips (*ρ_m_*), and all tips (*ρ_f_*), over described species. The summary trees of Amphibians, Seed plants and Squamates have nodes with more than two children; the share of branching events at them is in Results. *λ* ratio: speciation rate under the full tree’s fraction over speciation rate under the matching fraction, on the pruned tree; it equals *ρ_m_*/*ρ_f_* where the restriction does not bind and differs where extinction is forced to zero. *ε*: extinction fraction with 95% profile-likelihood interval. An interval whose estimate is zero starts at zero. †Median and range over the source’s taxon-complete posterior set and ‡over the set with dates drawn at random (trees fitted: Supplement S9), in place of the summary-tree estimate and its profile interval; *s*family fit on the summary tree. The summary-tree fits of these groups are in Supplement S9. LR: likelihood-ratio statistic of the matching fit against the fit with the full tree’s fraction (zero where the restriction does not bind); outcome on a summary tree “rescaled” (only *λ* and *μ* change) or “*μ* = 0” (extinction forced to zero), and over a set its median (range) and the trees in which extinction is forced to zero. A test is made only on a binary summary tree whose node ages come from a dating analysis (Fish, Mammals and Sharks): an asterisk marks a restriction rejected at the Bonferroni level 0.0167 (statistic above 4.53; 0 of the 3 tests; Materials and methods); the summary trees of Amphibians, Seed plants and Squamates and Birds are not tested (Materials and methods) and their statistics are reported without a test. *r*: net diversification rate (per Myr) on the full and the pruned tree. Family sampling: *ε* and *r* on the pruned tree with one sampling fraction per family (Materials and methods).

The full tree and the pruned tree, each with its own fraction, give net diversification rates with no common direction: over the archived tree sets, where the full trees are binary, the ratio of *r* on the full to *r* on the pruned tree ranges from 0.89 to 1.07, and on the summary trees *r* is higher on the full tree in 2 groups and lower in 5. The two fits share the sequenced tips, so their intervals are not independent and their overlap, in 3 groups, is not a test of a difference. The largest difference is in Seed plants, where *r* on the full tree is 1.30 times *r* on the pruned tree; 79.0% of the branching events of the full seed-plant tree, and 0.16% of the pruned tree’s, are at polytomies and its tips exceed its named species (Table 1), so this ratio reflects how tips and polytomies are counted more than branching times estimated from sequence data. The pruned seed-plant tree has an extinction fraction of 0.97, close to its upper limit of one, with a narrow interval; a constant-rate model that places extinction this close to speciation does not describe the tree, and the conditional profile interval does not measure how well the rates are estimated. With one sampling fraction per family instead of one per tree, the extinction fraction of the pruned tree is lower than under uniform sampling in 7 of the 7 groups (Supplement S9); the family likelihood is an approximation that changes with the order of children by more than some of the likelihood-ratio statistics above (Supplement S9), so these estimates are descriptive.

## Discussion

### Using a downloaded tree

The counts translate into a check a user can make before fitting a model. Take a squamate study: the largest tree (Tonini et al., 2016) has 12,252 described species as its denominator in Table 1, 0.56 of its tips are sequenced, and its source distributes a tree of its sequenced species, undated. A diversification analysis that should rest on branch lengths estimated from sequence data uses a dated tree of the sequenced species with a sampling fraction of 0.44 (sequenced tips over described species). The reference tree pruned to its sequenced tips is not such a tree: its node ages were estimated together with the taxonomically placed species, and as a consensus summary it keeps polytomies whose ages come from the summary. The squamate posterior tree set archived by Tonini et al. (2016), which also contains the taxonomically placed species, gives a squamate crown age of 186.4 Ma (177.0–203.1), an interval within the uniform calibration prior on the crown that spans 0.74 of its width (Supplement S6). For Amphibians, the frog tree of Portik et al. (2023) alone holds more sequenced species than the reference tree’s sequenced tips (Supplement S1), and with the salamander tree of Stewart and Wiens (2025) it covers frogs and salamanders, so a user who needs sequenced tips of these orders takes those trees rather than pruning the reference tree; caecilians remain only in the reference tree. Trait models are affected in the same way: on a tree completed by taxonomy, the branches to added species follow the placement algorithm, not the data, which biases trait-dependent diversification analyses (Rabosky, 2015).

A mismatched fraction moves *λ*, *μ* and *ε*, the quantities most often interpreted (Cusimano and Renner, 2010; Höhna et al., 2011), and moves *r* only when it forces *μ* to zero. I did not survey how published analyses of these trees set the sampling fraction, so the example shows the consequence of the error in each group, not how often it is made. Fitting the full tree is a different analysis, and the ratio of *r* on the full to *r* on the pruned tree has no common direction (Results); on the summary trees the lower values are those of the squamate and amphibian consensus trees (0.81 and 0.83) and the upper that of the seed-plant tree, with its polytomies (Supplement S9).

### Coverage in context

For every group in which the distinction matters, the source already distributes a tree of its sequenced species (Results), and the Open Tree of Life flags which tips come from an input phylogeny. What the count adds is a common basis across groups, with the molecular, dated and date-uncertainty levels counted the same way in each. On that basis a minority of the sequenced species of the reference trees are among the Open Tree’s phylogeny-placed species (Results).

### How date uncertainty is archived

An analysis integrates over dating uncertainty by fitting over a posterior or bootstrap set (de Villemereuil et al., 2012; Rangel et al., 2015). From the summary trees archived with node intervals (Cnidaria, Echinoderms, Gastropods and Sponges; Results) node ages can be drawn, but node ages are correlated and ordered, so drawing them independently from their intervals approximates the uncertainty one node at a time and gives no joint distribution of trees. The posterior sets of Amphibians, Cephalopods, Mammals, Squamates and Turtles carry the posterior of a Bayesian dating (Supplement S6). The bootstrap sets of Bryophytes and Diatoms are treePL datings in which the root sits on its maximum in every tree and the diatom crown median on a hard bound, so their variation is bootstrap variation under fixed or bounded calibrations (Supplement S6). The shark sets vary through root ages drawn from the calibration prior (Materials and methods), so their variation at every node carries that prior, and the taxon addition changed the relationships and ages of the sequenced species (Results); the source’s trees of its sequenced species, not the complete trees pruned, carry its dating of the sequence data. The largest-tree rule favours trees completed by taxonomy (Results); some groups have smaller trees archived with dating uncertainty, for example the posterior sets of bird trees restricted to sequenced species (Jetz et al., 2012), so a user who chooses a tree for its dates can reach higher coverage with date uncertainty available.

### Limitations

The reference trees come from a search of the sources listed in Materials and methods with no fixed query protocol, and from one tree per group, whose count under other partitions of the described species is in Supplement S7. Trees the search did not find can only add species to the 19.4% of described species named in any tree found; because each level is counted on the largest tree found, a larger tree replacing a reference tree can raise or lower the count at any level, and the other levels were not counted over all trees. The classification of tips is not a bound in the same sense: in 3 groups the sequenced tips are stated by the source and not listed, which affects 0.74% of molecular coverage. For several invertebrate groups the reference tree samples one subclade (echinoids for Echinoderms, Scleractinia for Cnidaria, Demospongiae for Sponges, decapods for Crustaceans, freshwater mussels for Bivalves) or is a phylogenomic backbone (Gastropods), and the search missed larger published trees of these and other groups, which are not counted: a Sanger-sequence tree of bivalves across the class (Combosch et al., 2017), a supertree of 1,547 scleractinian species built from a molecular tree of 474 species, morphological trees and a taxonomic tree, and dated by fitting the sequence data to the supertree topology with fossil calibrations (Huang and Roy, 2015), dated trees of true crabs (Wolfe et al., 2024) and of amphipods (Copilaş-Ciocianu et al., 2020), an exon-capture tree of brittle stars (O’Hara et al., 2017), a tree of cone snails (Puillandre et al., 2014), a small-subunit rRNA tree of demosponges (Redmond et al., 2013), a small-subunit rRNA tree of nematodes (van Megen et al., 2009), and the dated tree of Agaricomycetes of Varga et al. (2019), which is larger than the fungal reference tree; phylogenomic trees of flatworms (Laumer et al., 2015) and annelids (Struck et al., 2011) would give those groups a reference tree. A smaller tree can also place more species with sequence data than the reference tree of its group, as the frog tree of Portik et al. (2023) does for Amphibians. The count rests on one tree, that of seed plants (Results, Coverage by group). The denominators come from group-level authorities of different kinds; Table 2 gives each level with a single Catalogue of Life count. A sequenced tip may rest on one locus, so molecular coverage counts species placed by sequence data, not species with well-estimated branch lengths. The fits of the worked example are conditional on a constant-rate model and on uniform or family-level sampling, on one summary tree per group and, where the source archives them, on a sample of its dated trees.

## Conclusions

A source archive that lets a user recompute the levels counted here holds the tree, the list of tips in the molecular analysis or a tree of them, the posterior or bootstrap tree set or a summary tree with node intervals, and the calibration file; a run sampling from the prior shows how far the data moved the dates. Among the groups with the highest named coverage, the archives of the largest trees of Seed plants and Fish carry no date uncertainty for their sequenced species, and those of Birds and Sharks carry sets whose dates are drawn from published studies or under sampled root ages; an archived posterior set of dated trees for Seed plants, Fish, Birds or Sharks would extend date uncertainty available to the sequenced species of Seed plants and Fish, more than any other addition to the trees counted here, and give those of Birds and Sharks the posterior of a dating of their sequence data.

## Data availability

The trees whose source licences allow redistribution (255 of the 263; the other 8 are listed with the source of each), the per-group table, the molecular tip lists and the crown-age and calibration registers are in data release v1.1.5 of https://github.com/franciscorichter/phylo-species-atlas, tagged v1.1.5 in that repository and archived on Zenodo (version DOI: https://doi.org/10.5281/zenodo.24088448) under the concept DOI https://doi.org/10.5281/zenodo.20127157, which resolves to every version. The source trees remain in their original archives, as cited.

## Code availability

The code that rebuilds every derived file and the script that recomputes every reported number are in the same release; Supplement S10 lists them.

## Acknowledgements

I thank the authors of the phylogenetic studies used here for archiving their trees, alignments and dated tree sets.

## Funding

This research was supported through the research programme “Statistical Frontiers in Dynamic Network Modeling”, funded by the Swiss National Science Foundation (project FNS 240065), directed by Prof. Ernst-Jan Camiel Wit.

## Author contributions and use of AI

Since February 2026 this work has been carried out with large language models from Anthropic (Claude, used through the Claude Code command-line tool; Claude Opus 5.5 and Claude Fable 5.1 and 5.2 (model identifiers claude-opus-5-5, claude-fable-5-1 and claude-fable-5-2) for this version and earlier Claude Opus models before it). The models wrote and ran the code for data retrieval, name resolution, analysis and figures; searched the literature and the data archives for phylogenies and their documentation; and drafted text, which I edited. Each reference was checked against its DOI record.

The article, the data and the code were checked by further model runs separate from the ones that produced them; each error they reported was recorded, and each correction was checked by a run other than the one that made it. Analytical choices follow rules written before they were applied. Every number in the article and its Supplement is generated by code from archived files, and a separate script recomputes each one (Code availability). The agreement of the two shows that the numbers follow from the stated rules, not that the rules are correct: the branch lengths corrected in data releases v1.0.3 to v1.0.8 were found by checking the files against their sources (Supplement S10).

In the continuous procedure (Materials and methods, Continuous data collection), code written by the models screens new deposits and the models check each candidate against its source study; I approve each data release before it is published.

## Competing interests

The author declares no conflict of interest.

## Notes

https://franciscorichter.github.io/phylo-species-atlas

https://doi.org/10.5281/zenodo.24088448

https://github.com/franciscorichter/phylo-species-atlas

### Competing Interest Statement

The authors have declared no competing interest.

### Summary of Updates

Each group's described species count is now the figure its cited source states, taken from the authority's release nearest the Catalogue of Life release the names are matched to (COL26.9) or from the publication cited; the denominator becomes 2,059,063 described eukaryotic species. The headline counts species named in any tree of the catalogue (19.4%); the reference tree of each group places 7.6% with molecular data and 5.5% in a dated tree, and date uncertainty is available for 1.2%. A new table in Supplement S7 recounts coverage with one tree per group when the groups are the Catalogue of Life classes, orders, families or phyla: with classes, orders or families the count is within 1.1% of the count over every tree. The worked example is refitted on the new sampling fractions; extinction is forced to zero in 100 of 100 squamate and 89 of 100 amphibian posterior trees. Three figures show the Species Phylogenies Catalogue, the online platform of the data: the procedure that keeps it current (Fig. 1), the groups around the tree of life with the share named in a reference tree (Fig. 2), and the four levels drawn as the site draws them, with described species against the estimated number of species (Fig. 3, replacing the bar chart of the levels); the per-group bar chart is removed, since the ring, the scatter of named coverage against molecular fraction and the crown-age figure carry its content, and all figures share one palette in which each colour has one meaning. The use-of-AI statement states what the language models did, how their work was checked by separate model runs and a script that recomputes every number, that the author approves each data release, and names the models used for this version. The text is shortened: each fact is stated in one place (Methods define, Results state, the Discussion interprets, captions explain the display), in the article and in the Supplement. Data release v1.1.5 holds the trees whose source licences allow redistribution, the registers and the code, including the scripts of the six figures.

