## Supplementary Information (Supplements S1 to S10) for "Described eukaryotic species in published species-level phylogenies: placement by molecular data, dates and date uncertainty"

doi: 10.64898/2026.05.31.729127
Revised: November 2026
Data: release v1.1.5

**Francisco Richter** ORCID 0000-0002-0924-4613
Università della Svizzera italiana (USI), Lugano, Switzerland
✉

### Contents

**Supplement S1: groups, described species and reference trees** 1
**Supplement S2: tips removed and merged** 5
**Supplement S3: name resolution** 6
**Supplement S4: molecular tips** 7
**Supplement S5: time scale of the reference trees** 8
**Supplement S6: crown ages and archived intervals** 9
**Supplement S7: sensitivity analyses** 11
**Supplement S8: comparison with the Open Tree of Life** 13
**Supplement S9: worked example** 13
**Supplement S10: data and code** 17

### Supplement S1: groups, described species and reference trees

The 60 groups of eukaryotes (Materials and methods of the article, Groups and reference trees) and their described species are listed in Table S1, with the source of each count. The fish count is that of the ray-finned fishes (Cladistii and Actinopteri) in Eschmeyer's Catalog of Fishes; hagfishes, lampreys, cartilaginous fishes, coelacanths and lungfishes are counted in their own groups, so the counts do not overlap. The insect count makes up 45.2% of the sum of the counts; Supplement S7 varies it, and replaces every count by a Catalogue of Life count.

The seed-plant reference tree is the tree of all species of Smith and Brown (2018), and its molecular tips are those of the tree of the species with GenBank sequences of the same study, both on an Open Tree backbone. NDexp marks the node-dated mammal trees of Upham et al. (2019), whose calibrations carry exponential priors.

Table S1 also lists every tree of the catalogue: 263 files, of which the 218 family-level trees of one compilation are shown as one row. Its last block lists the 36 groups for which the search found no downloadable species-level tree, which hold 219,661 described species.

Table S2 gives, for each of the 24 reference trees, the tree used, the archive it comes from and the rule of Materials and methods of the article (Groups and reference trees) by which it is chosen among the files of its source. 15 trees are used as their source archives them (A) and 3 are named releases of trees that are updated continuously (R); the others are a study's chronogram (C: Bryophytes, Gastropods and Turtles), the tree of a study with more species carrying usable branch lengths (B: Diatoms), the first tree of a set (F: Fish) and the maximum-clade-credibility tree of a set (M: Sharks). The shark source archives the sets of two dating scenarios, with one and with ten calibration fossils; its Supplementary Information states that it summarises its results from the ten-fossil trees, and that set is used here. Both scenarios draw their root ages from the same calibration density, and the one-fossil set gives the same crown interval to within a fraction of a million years; the root ages the source reports for both are the range of root heights over each set. For Birds, the version of release AvesData 1.3 used is chosen by the rule for a synthesis aligned to several versions of one taxonomy (Materials and methods of the article, Groups and reference trees); the tips of the Clements 2023 version that an input phylogeny places are counted on its tree of phylogeny-placed tips (McTavish et al., 2025), and that version is listed in Table 1 of the main text. The data release holds a copy of each source file and the evidence for each choice, and its acceptance test checks that the branch lengths of each reference tree's archived source equal those of that file.

**Table S1 | Groups, described species and the trees of the catalogue.** Each row is one tree of the catalogue (the data release redistributes all but the 8 whose sources state no licence: cephalopods, frogs, gastropods, insects, primates, seed plants, sharks and sponges), the family-level trees of one compilation (one row), a cited dataset that is not redistributed, or a group with no tree. Bold: the reference tree of a group. Tips: distinct tip labels of the file. Names: distinct Catalogue of Life accepted species among the tips. Desc.: described species; for a reference tree the group's count, for another tree the count for its clade where a source for it is recorded. %: Names over Desc. Dated, for a reference tree: Yes, a chronogram in absolute time; Rel., ultrametric in relative time; No, not ultrametric; for another tree: Ultr., ultrametric (root-to-tip coefficient of variation at most 0.05), whose time scale is not checked; No, not ultrametric. [b]Genus-level tips. [c]Genome Taxonomy Database species clusters, the unit of both numerator and denominator for prokaryotes, which never enter a eukaryotic total. [d]Cited and not redistributed, at the request of the TimeTree project. [e]Preprint. [f]Relative time. [g]Ultrametric. Abbreviations: MPE, Mol. Phylogenet. Evol.; MBE, Mol. Biol. Evol.; NAR, Nucleic Acids Res.; CoL, Catalogue of Life; WoRMS, World Register of Marine Species.

| Group | Study | Year | Journal | Tips | Names | Desc. | % | Dated | Source of Desc. |
|---|---|---|---|---|---|---|---|---|---|
| ***Vertebrates*** | | | | | | | | | |
| **Fish** | Rabosky et al. (2018) | 2018 | Nature | 31,347 | 30,973 | 36,288 | 85.4 | Yes | Eschmeyer's Catalog of Fishes (21 Sep. 2026) |
| **Birds** | McTavish et al. (2025) | 2025 | PNAS | 10,824 | 10,469 | 11,185 | 93.6 | Yes | IUCN Red List 2026-1, Table 1a (HBW-BirdLife v10) |
| **Squamates** | Tonini et al. (2016) | 2016 | Biol Conserv | 9,754 | 8,953 | 12,252 | 73.1 | Yes | Reptile Database, June 2026 release (Uetz et al.) |
| **Amphibians** | Jetz and Pyron (2018) | 2018 | Nat Ecol Evol | 7,238 | 6,947 | 9,088 | 76.4 | Yes | AmphibiaWeb (11 Sep. 2026) |
| **Mammals** | Upham et al. (2019) | 2019 | PLOS Biol | 5,911 | 5,625 | 6,887 | 81.7 | Yes | Mammal Diversity Database v2.5 (2026) |
| **Sharks** | Stein et al. (2018) | 2018 | Nat Ecol Evol | 1,192 | 1,150 | 1,331 | 86.4 | Yes | Eschmeyer's Catalog of Fishes (21 Sep. 2026) |
| **Turtles** | Thomson et al. (2021) | 2021 | PNAS | 283 | 256 | 359 | 71.3 | Yes | Turtle Taxonomy Working Group 2025 (10th ed.), living species |
| **Crocodilians** | Condamine et al. (2019) | 2019 | Ecol Lett | 25 | 20 | 27 | 74.1 | Yes | The Reptile Database, June 2026 (via COL26.9) |
| Frogs | Portik et al. (2023) | 2023 | MPE | 5,326 | 5,235 | 7,486 | 69.9 | Ultr.[g] | AmphibiaWeb |
| Neotrop. fish | Tagliacollo et al. (2024) | 2024 | Front Bioinf | 3,167 | 3,052 | — | — | Ultr.[g] | — |
| Salamanders | Stewart and Wiens (2025) | 2025 | MPE | 796 | 789 | 814 | 96.9 | Ultr.[g] | AmphibiaWeb |
| Parrots | Burgio et al. (2019) | 2019 | Data Brief | 413 | 388 | 413 | 93.9 | Ultr.[g] | IUCN |
| Primates | Arnold et al. (2010) | 2010 | Evol Anthro | 301 | 262 | 522 | 50.2 | Ultr.[g] | IUCN |
| Carnivora | Nyakatura and Bininda-Emonds (2012) | 2012 | BMC Biol | 294 | 286 | 294 | 97.3 | Ultr.[g] | IUCN |
| Cetaceans | McGowen et al. (2020) | 2020 | Syst Biol | 85 | 84 | 92 | 91.3 | Ultr.[g] | WoRMS |
| ***Plants*** | | | | | | | | | |
| **Seed plants** | Smith and Brown (2018) | 2018 | Am J Bot | 348,886 | 279,049 | 354,303 | 78.8 | Yes | World Flora Online, as in COL26.9 (2026-09-11) |

| Group | Study | Year | Journal | Tips | Names | Desc. | % | Dated | Source of Desc. |
|---|---|---|---|---|---|---|---|---|---|
| **Ferns** | Nitta et al. (2022) | 2022 | Front Plant Sci | 6,216 | 6,097 | 11,999 | 50.8 | Yes | PPG II, pteridogroup/ppg v2.0.0 (15 Sep. 2026) |
| **Bryophytes** | Bechteler et al. (2023) | 2023 | Am J Bot | 533 | 505 | 19,273 | 2.6 | Yes | Catalogue of Life COL26.9 |
| Orchids | Thompson et al. (2023) | 2023 | PNAS | 1,475 | 1,433 | 28,000 | 5.1 | Ultr.[g] | CoL |
| Grasses | Grass Phylogeny Working Group III (2025) | 2025 | New Phytol | 1,153 | 986 | 12,000 | 8.2 | No | CoL |
| Solanaceae | Särkinen et al. (2013) | 2013 | BMC Evol Biol | 1,076 | 0 | — | — | Ultr.[g] | — |
| Cacti | Thompson et al. (2026) | 2026 | Sci Data | 1,063 | 1,014 | 1,900 | 53.4 | Ultr.[g] | CoL |
| Eucalyptus | Crisp et al. (2024) | 2024 | J Syst Evol | 399 | 103 | — | — | No | — |
| Asteraceae | Mandel et al. (2019) | 2019 | PNAS | 256 | 210 | 34,000 | 0.6 | No | CoL |
| Pinaceae (in Seed plants) | Ran et al. (2018) | 2018 | MPE | 16[b] | 0 | — | — | Ultr.[g] | — |
| ***Arthropods*** | | | | | | | | | |
| **Insects** | Chesters et al. (2023) | 2023 | Mol Ecol Res | 53,373 | 42,078 | 929,676 | 4.5 | Rel.[f] | Catalogue of Life COL26.9 |
| **Spiders** | Macías-Hernández et al. (2020) | 2020 | Diversity | 1,431 | 888 | 54,244 | 1.6 | Yes | World Spider Catalog v27 (read 4 Nov. 2026) |
| **Crustaceans** | Wolfe et al. (2019) | 2019 | Proc R Soc B | 95 | 65 | 67,735 | 0.1 | No | Zhang 2013 (Zootaxa 3703) |
| Bees | Henríquez-Piskulich et al. (2024) | 2024 | MPE | 4,586 | 4,369 | 20,507 | 21.3 | Ultr.[g] | Discover Life |
| Butterflies | Kawahara et al. (2023) | 2023 | Nat Ecol Evol | 2,258 | 1,128 | 19,000 | 5.9 | Ultr.[g] | CoL |
| Hemiptera | Johnson et al. (2018) | 2018 | PNAS | 192 | 147 | — | — | No | — |
| Termites | Hellemans et al. (2024) | 2024 | Nat Comms | 138 | 0 | — | — | Ultr.[g] | — |
| Ants | Romiguier et al. (2022) | 2022 | Curr Biol | 83 | 62 | 14,415 | 0.4 | No | AntCat |
| ***Other animals*** | | | | | | | | | |
| **Bryozoa** | Orr et al. (2022) | 2022 | Sci Adv | 565 | 337 | 6,613 | 5.1 | No | WoRMS (accessed 4 Nov. 2026) |
| **Cnidaria** | Vaga et al. (2025) | 2025 | Nature | 289 | 258 | 11,597 | 2.2 | Yes | WoRMS (accessed 4 Nov. 2026) |
| **Nematodes** | Qing et al. (2025) | 2025 | Syst Biol | 156 | 101 | 28,415 | 0.4 | No | Hodda 2022a (28,537 species including at least 122 fossil) |
| **Sponges** | Lavrov et al. (2023) | 2023 | PLOS ONE | 119 | 86 | 9,864 | 0.9 | Yes | World Porifera Database (15 Aug. 2026) |
| **Cephalopods** | Basava et al. (2024) | 2024 | bioRxiv[e] | 81 | 80 | 868 | 9.2 | Yes | MolluscaBase, in Catalogue of Life COL26.9 |
| **Bivalves** | Pfeiffer et al. (2019) | 2019 | MPE | 73 | 44 | 10,043 | 0.4 | No | MolluscaBase, in Catalogue of Life COL26.9 |
| **Echinoderms** | Mongiardino Koch et al. (2022) | 2022 | eLife | 65 | 62 | 7,749 | 0.8 | Yes | WoRMS (accessed 4 Nov. 2026) |
| **Gastropods** | Zapata et al. (2014) | 2014 | Proc R Soc B | 40 | 39 | 76,288 | 0.1 | Yes | MolluscaBase, in Catalogue of Life COL26.9 |
| ***Microbes & protists*** | | | | | | | | | |
| **Bacteria** | Parks et al. (2026) | 2026 | NAR | 136,646 | 136,646[c] | 136,646[c] | 100.0 | No | GTDB R10-RS226 species clusters |
| **Archaea** | Parks et al. (2026) | 2026 | NAR | 6,968 | 6,968[c] | 6,968[c] | 100.0 | No | GTDB R10-RS226 species clusters |
| **Fungi** | Li et al. (2021) | 2021 | Curr Biol | 1,620 | 1,396 | 156,891 | 0.9 | No | Catalogue of Life COL26.9 (Species Fungorum Plus; Microsporidia) |
| **Diatoms** | Nakov et al. (2018) | 2018 | New Phytol | 1,113 | 39 | 16,427 | 0.2 | Yes | AlgaeBase (Guiry 2024) |
| ***Cross-cutting*** | | | | | | | | | |
| Eukaryote root | Williamson et al. (2025) | 2025 | Nature | 100 | 0 | — | — | No | — |
| Tetrapod families | Condamine et al. (2019) | 2019 | Ecol Lett | 218 trees | — | — | — | 218 of 218 | — |
| TimeTree | Hedges et al. (2015) | 2015 | MBE | — | — | — | — | —[d] | — |
| ***Not yet represented*** | | | | | | | | | |
| Acari | — | — | — | 0 | 0 | 54,908 | 0.0 | — | Zhang 2013 (Zootaxa 3703), extant species |
| Flatworms | — | — | — | 0 | 0 | 29,487 | 0.0 | — | Zhang 2013 (29,488 species including 1 fossil) |
| Annelids | — | — | — | 0 | 0 | 17,340 | 0.0 | — | Catalogue of Life COL26.9 |
| Myriapods | — | — | — | 0 | 0 | 16,072 | 0.0 | — | Chapman 2009 |

| Group | Study | Year | Journal | Tips | Names | Desc. | % | Dated | Source of Desc. |
|---|---|---|---|---|---|---|---|---|---|
| Other arachnids | — | — | — | 0 | 0 | 12,765 | 0.0 | — | Catalogue of Life COL26.9 |
| Green algae | — | — | — | 0 | 0 | 11,791 | 0.0 | — | AlgaeBase via Guiry 2024 (counts to Sep. 2023), extant |
| Other hexapods | — | — | — | 0 | 0 | 10,138 | 0.0 | — | Catalogue of Life COL26.9 |
| Foraminifera | — | — | — | 0 | 0 | 8,972 | 0.0 | — | World Foraminifera Database / WoRMS (4 Nov. 2026) |
| Rhodophyta | — | — | — | 0 | 0 | 7,276 | 0.0 | — | AlgaeBase via Guiry 2024 (1 Nov. 2023), living species |
| Apicomplexa | — | — | — | 0 | 0 | 6,000 | 0.0 | — | Morrison 2008 |
| Ciliates | — | — | — | 0 | 0 | 4,500 | 0.0 | — | Foissner et al. 2008 |
| Other phyla | — | — | — | 0 | 0 | 3,863 | 0.0 | — | Catalogue of Life COL26.9 |
| Other protists | — | — | — | 0 | 0 | 3,353 | 0.0 | — | Pawlowski et al. 2012 (PLoS Biol 10: e1001419), Table S1: 29 lineages in no other group |
| Tunicates | — | — | — | 0 | 0 | 3,020 | 0.0 | — | Appeltans et al. 2012 |
| Myxozoa | — | — | — | 0 | 0 | 3,012 | 0.0 | — | World List of Myxozoa (WoRMS), in Catalogue of Life COL26.9 |
| Scorpiones | — | — | — | 0 | 0 | 2,954 | 0.0 | — | The Scorpion Files (Rein; Wayback 3 Oct. 2026) |
| Amoebozoa | — | — | — | 0 | 0 | 2,400 | 0.0 | — | Pawlowski et al. 2012 |
| Dinoflagellates | — | — | — | 0 | 0 | 2,377 | 0.0 | — | Gómez 2012 checklist (CICIMAR Oceánides 27: 65-140) |
| Other Mollusca | — | — | — | 0 | 0 | 2,199 | 0.0 | — | MolluscaBase, in Catalogue of Life COL26.9 |
| Brown algae | — | — | — | 0 | 0 | 2,094 | 0.0 | — | AlgaeBase (Guiry 2024) |
| Rotifera | — | — | — | 0 | 0 | 2,030 | 0.0 | — | Segers 2007 (about 2,030) |
| Oomycetes | — | — | — | 0 | 0 | 1,673 | 0.0 | — | Catalogue of Life COL26.9 (Species Fungorum Plus) |
| Euglenozoa | — | — | — | 0 | 0 | 1,520 | 0.0 | — | CBOL Protist Working Group (Pawlowski et al. 2012), Table S1 |
| Tardigrada | — | — | — | 0 | 0 | 1,505 | 0.0 | — | Degma & Guidetti, Actual checklist of Tardigrada species (2025) |
| Lycophytes | — | — | — | 0 | 0 | 1,467 | 0.0 | — | PPG II, pteridogroup/ppg v2.0.0 (15 Sep. 2026) |
| Pycnogonida | — | — | — | 0 | 0 | 1,431 | 0.0 | — | WoRMS (Pycnobase) 2026-10-01 |
| Acanthocephala | — | — | — | 0 | 0 | 1,325 | 0.0 | — | Catalogue of Life COL26.9 |
| Nemerteans | — | — | — | 0 | 0 | 1,275 | 0.0 | — | Kajihara et al. 2008 |
| Chrysophyta | — | — | — | 0 | 0 | 1,216 | 0.0 | — | AlgaeBase (Guiry 2024) |
| Cercozoa | — | — | — | 0 | 0 | 600 | 0.0 | — | Pawlowski et al. 2012 (CBOL ProWG, Table S1) |
| Haptophytes | — | — | — | 0 | 0 | 517 | 0.0 | — | AlgaeBase (Guiry 2024; living species, 1 Nov. 2023) |
| Choanoflagellates | — | — | — | 0 | 0 | 244 | 0.0 | — | Carr et al. 2008 ('over 244') |
| Ctenophores | — | — | — | 0 | 0 | 186 | 0.0 | — | WoRMS (accessed 4 Nov. 2026) |
| Hagfish | — | — | — | 0 | 0 | 93 | 0.0 | — | Eschmeyer's Catalog of Fishes (20 Oct. 2026) |
| Lampreys | — | — | — | 0 | 0 | 49 | 0.0 | — | Eschmeyer's Catalog of Fishes (21 Sep. 2026) |
| Other vertebrates | — | — | — | 0 | 0 | 9 | 0.0 | — | The Reptile Database (Jun. 2026) and Eschmeyer's Catalog of Fishes (21 Sep. 2026) |

**Table S2 | Provenance of the reference trees.** Tree used: the tree of the source archive. Rule: A, the tree as the source archives it; R, a named release of a tree updated continuously; C, the study's chronogram, which covers all tips of its other tree; B, of two trees of the study, the one with more species carrying usable branch lengths; F, the first tree of a set with the same tips, of which the source designates none; M, the maximum-clade-credibility tree of a set of dated trees.

| Group | Source | Tree used | Rule | Archive |
|---|---|---|---|---|
| *Vertebrates* | | | | |
| Amphibians | Jetz and Pyron (2018) | consensus chronogram | A | https://doi.org/10.5061/dryad.cc3n6j5 |
| Birds | McTavish et al. (2025) | Aves 1.3, Clements 2021 | R | https://github.com/McTavishLab/AvesData |
| Crocodilians | Condamine et al. (2019) | Crocodylia tree | A | https://doi.org/10.6084/m9.figshare.9555842 |
| Fish | Rabosky et al. (2018) | tree 1 | F | https://fishtreeoflife.org |
| Mammals | Upham et al. (2019) | NDexp MCC chronogram | A | https://doi.org/10.5061/dryad.tb03d03 |
| Sharks | Stein et al. (2018) | MCC tree | M | https://data.vertlife.org/sharktree |
| Squamates | Tonini et al. (2016) | all-compatible consensus chronogram | A | https://doi.org/10.5061/dryad.db005 |
| Turtles | Thomson et al. (2021) | MCC chronogram | C | https://doi.org/10.5061/dryad.jh9w0vt8w |
| *Plants* | | | | |
| Bryophytes | Bechteler et al. (2023) | treePL chronogram | C | https://doi.org/10.5061/dryad.3j9kd51qm |
| Ferns | Nitta et al. (2022) | release v1.8.0 | R | https://github.com/fernphy/ftol_data |
| Seed plants | Smith and Brown (2018) | treePL chronogram | A | https://github.com/FePhyFoFum/big_seed_plant_trees |
| *Arthropods* | | | | |
| Crustaceans | Wolfe et al. (2019) | PhyloBayes CAT-GTR tree | A | https://doi.org/10.5061/dryad.k7505mn |
| Insects | Chesters et al. (2023) | Insecta_v2 | R | https://insectphylo.org |
| Spiders | Macías-Hernández et al. (2020) | preferred chronogram | A | https://doi.org/10.5281/zenodo.3941712 |
| *Other animals* | | | | |
| Bivalves | Pfeiffer et al. (2019) | ASTRAL species tree | A | https://doi.org/10.5061/dryad.8575jr5 |
| Bryozoa | Orr et al. (2022) | trimmed ML tree | A | https://doi.org/10.5061/dryad.2v6wwpzp9 |
| Cephalopods | Basava et al. (2024) | consensus chronogram | A | https://github.com/kcbasava/ceph-brain-evolution |
| Cnidaria | Vaga et al. (2025) | MCC chronogram | A | https://doi.org/10.6084/m9.figshare.29242487 |
| Echinoderms | Mongiardino Koch et al. (2022) | PhyloBayes chronogram | A | https://doi.org/10.5061/dryad.brv15dv9t |
| Gastropods | Zapata et al. (2014) | MCMCTree chronogram | C | https://bitbucket.org/caseywdunn/gastropoda |
| Nematodes | Qing et al. (2025) | ML tree of the 50% amino-acid matrix | A | https://doi.org/10.5061/dryad.b2rbnzsnt |
| Sponges | Lavrov et al. (2023) | PhyloBayes chronogram | A | https://github.com/LavrovLab/Demosponge-phylogeny-supplementary-data |
| *Microbes & protists* | | | | |
| Diatoms | Nakov et al. (2018) | tree 1: treePL-dated best ML tree | B | https://doi.org/10.5281/zenodo.583628 |
| Fungi | Li et al. (2021) | species tree as archived | A | https://doi.org/10.6084/m9.figshare.12751736 |

### Supplement S2: tips removed and merged

Each reference tree is built from its archived source by rule (Table S3). First, tips outside the group (outgroups and misplaced taxa) are removed: 621 tips from 13 trees, each listed with its reason in the data release. Then, reading the tips from left to right, a tip whose species key has already been seen is removed, so that one tip per species remains: 7,532 tips from 8 trees. The species key of a label is its first pair of a genus and an epithet. A capitalised token that ends in a suffix of a family or higher rank (for example -idae or -aceae) is read as a genus only when the Catalogue of Life lists a genus of that name (338 such genera in COL26.9). A label without a key is never merged with another tip: labels in open nomenclature (cf., aff., sp.), hybrids, and labels that are codes or sample identifiers. These are 12,670 of the 481,229 tips. The counts are those of the files as distributed; the fish all-taxon trees distributed by FishTreeOfLife hold slightly fewer tips than the total the source paper states. Finally, nodes left with one child are removed and their branch lengths summed, and a root left with one child is replaced by that child, so the root of a reference tree is the most recent common ancestor of its tips.

**Table S3 | Tips removed and merged in building each reference tree.** Source: tips of the archived source tree. Outside group: tips removed as outgroups or misplaced taxa. Merged: tips removed because an earlier tip has the same species key. No key: tips of the reference tree whose label has no species key, which are never merged.

| Group | Source | Outside group | Merged | Tips | No key |
|---|---|---|---|---|---|
| *Vertebrates* | | | | | |
| Amphibians | 7,239 | 1 | 0 | 7,238 | 0 |
| Birds | 10,824 | 0 | 0 | 10,824 | 0 |
| Crocodilians | 25 | 0 | 0 | 25 | 25 |
| Fish | 31,516 | 0 | 169 | 31,347 | 10 |
| Mammals | 5,912 | 1 | 0 | 5,911 | 0 |
| Sharks | 1,192 | 0 | 0 | 1,192 | 2 |
| Squamates | 9,755 | 1 | 0 | 9,754 | 0 |
| Turtles | 288 | 2 | 3 | 283 | 4 |
| *Plants* | | | | | |
| Bryophytes | 533 | 0 | 0 | 533 | 9 |
| Ferns | 6,234 | 18 | 0 | 6,216 | 0 |
| Seed plants | 356,305 | 282 | 7,137 | 348,886 | 11,264 |
| *Arthropods* | | | | | |
| Crustaceans | 95 | 0 | 0 | 95 | 95 |
| Insects | 53,596 | 223 | 0 | 53,373 | 8 |
| Spiders | 1,456 | 0 | 25 | 1,431 | 506 |
| *Other animals* | | | | | |
| Bivalves | 73 | 0 | 0 | 73 | 23 |
| Bryozoa | 720 | 20 | 135 | 565 | 193 |
| Cephalopods | 81 | 0 | 0 | 81 | 0 |
| Cnidaria | 289 | 0 | 0 | 289 | 14 |
| Echinoderms | 66 | 1 | 0 | 65 | 1 |
| Gastropods | 54 | 14 | 0 | 40 | 1 |
| Nematodes | 166 | 10 | 0 | 156 | 25 |
| Sponges | 120 | 0 | 1 | 119 | 24 |
| *Microbes & protists* | | | | | |
| Diatoms | 1,171 | 20 | 38 | 1,113 | 341 |
| Fungi | 1,672 | 28 | 24 | 1,620 | 125 |

### Supplement S3: name resolution

Each tip label is resolved against the Catalogue of Life base release COL26.9 (ChecklistBank dataset 316321). The query is the label's species key (Supplement S2). It resolves to an accepted or provisionally accepted species with that genus and epithet; where there are several in different kingdoms, to the one in the group's kingdom, which is the kingdom of most of the group's unambiguous matches. Otherwise it resolves through a synonym to the accepted species the synonym points to, dropping targets in other kingdoms; a label with more than one remaining target is ambiguous and does not resolve. A comparison that ignores hyphens and case is tried when these find nothing. Two registers correct single labels, each row with its evidence: 97 labels that carry no binomial, such as lineage codes, are queried by the binomial the source's own table of codes gives (a label without a species key resolves only in this way). 86 of them resolve, the number per group given in Table S4, and 11 match no Catalogue species. In the other register, targets that name a different organism from the one the label names are dropped (1 in all).

Of the 481,229 tip labels, 59,239 resolve through a synonym and 55,193 (11.5%) do not resolve

(Table S4). A named species is a distinct accepted species among the resolved labels; 30,519 resolved labels reach an accepted species that another label of the same tree already reaches. No accepted species is reached from two groups. As a sensitivity, the 38,276 labels that carry a species key and match no Catalogue name were queried against the GBIF backbone taxonomy on 7–8 October 2026 (strict match, species rank); 31,947 of them match, adding 27,834 species not already counted. Labels without a species key, labels that match several accepted species, and labels that match only species of another kingdom are not queried.

**Table S4 | Resolution of tip labels to Catalogue of Life accepted species, by group.** Labels: tips of the reference tree. Accepted: labels matching an accepted or provisionally accepted species. Synonym: labels resolving through a synonym. Unresolved: labels that are not species labels, are ambiguous or are unmatched, and their percentage of the labels. Override: resolved labels queried by the binomial of the override register. Merged: resolved labels whose accepted species another label already reaches. Named: distinct accepted species, the named-coverage numerator. GBIF: species added when unmatched labels are resolved with the GBIF backbone.

| Group | Labels | Accepted | Synonym | Unresolved | % | Override | Merged | Named | GBIF |
|---|---|---|---|---|---|---|---|---|---|
| *Vertebrates* | | | | | | | | | |
| Amphibians | 7,238 | 5,903 | 1,296 | 39 | 0.5 | 0 | 252 | 6,947 | 18 |
| Birds | 10,824 | 10,107 | 403 | 314 | 2.9 | 0 | 41 | 10,469 | 286 |
| Crocodilians | 25 | 20 | 0 | 5 | 20.0 | 20 | 0 | 20 | 0 |
| Fish | 31,347 | 29,015 | 2,241 | 91 | 0.3 | 0 | 283 | 30,973 | 44 |
| Mammals | 5,911 | 5,309 | 430 | 172 | 2.9 | 0 | 114 | 5,625 | 147 |
| Sharks | 1,192 | 1,001 | 179 | 12 | 1.0 | 0 | 30 | 1,150 | 4 |
| Squamates | 9,754 | 8,710 | 372 | 672 | 6.9 | 1 | 129 | 8,953 | 598 |
| Turtles | 283 | 253 | 4 | 26 | 9.2 | 0 | 1 | 256 | 16 |
| *Plants* | | | | | | | | | |
| Bryophytes | 533 | 429 | 78 | 26 | 4.9 | 0 | 2 | 505 | 4 |
| Ferns | 6,216 | 6,045 | 71 | 100 | 1.6 | 0 | 19 | 6,097 | 93 |
| Seed plants | 348,886 | 254,985 | 53,469 | 40,432 | 11.6 | 0 | 29,405 | 279,049 | 16,873 |
| *Arthropods* | | | | | | | | | |
| Crustaceans | 95 | 61 | 4 | 30 | 31.6 | 65 | 0 | 65 | 3 |
| Insects | 53,373 | 41,825 | 474 | 11,074 | 20.7 | 0 | 221 | 42,078 | 8,930 |
| Spiders | 1,431 | 883 | 5 | 543 | 37.9 | 0 | 0 | 888 | 29 |
| *Other animals* | | | | | | | | | |
| Bivalves | 73 | 32 | 13 | 28 | 38.4 | 0 | 1 | 44 | 0 |
| Bryozoa | 565 | 331 | 6 | 228 | 40.4 | 0 | 0 | 337 | 27 |
| Cephalopods | 81 | 75 | 5 | 1 | 1.2 | 0 | 0 | 80 | 1 |
| Cnidaria | 289 | 253 | 6 | 30 | 10.4 | 0 | 1 | 258 | 3 |
| Echinoderms | 65 | 60 | 2 | 3 | 4.6 | 0 | 0 | 62 | 0 |
| Gastropods | 40 | 37 | 2 | 1 | 2.5 | 0 | 0 | 39 | 0 |
| Nematodes | 156 | 98 | 3 | 55 | 35.3 | 0 | 0 | 101 | 23 |
| Sponges | 119 | 81 | 6 | 32 | 26.9 | 0 | 1 | 86 | 5 |
| *Microbes & protists* | | | | | | | | | |
| Diatoms | 1,113 | 38 | 2 | 1,073 | 96.4 | 0 | 1 | 39 | 682 |
| Fungi | 1,620 | 1,246 | 168 | 206 | 12.7 | 0 | 18 | 1,396 | 48 |

### Supplement S4: molecular tips

The molecular tips of a reference tree, the sequenced tips of the article, are found by one of four routes (Table S5). T: the tips of a tree that the source built from its sequenced species (7 groups). L: the taxa of the source's sequence alignment, archived with the data release (11 groups). E: per-tip evidence from the source's own records, listed tip by tip in the data release (3 groups). S: the source states that every tip was sequenced (3 groups). For T and L, a tip of the reference tree is molecular when its species key equals a species key of the source tree or alignment, after both sets of labels are resolved through the source's own names map; a label without a key is compared as it is. For route T every source species is in the reference tree except the outgroups removed from it, so the molecular tips are the source's labels less those that share a species key with another label, which the reference tree carries as one tip (columns Source labels, Merged and Unmatched). Matching the labels exactly gives the column Exact. It is lower where the source labels carry specimen codes or another spelling, and zero for Diatoms and Mammals, whose source labels are written in another format; it is not used. The molecular species are the distinct accepted species among the molecular tips. The product of named species and molecular fraction, a sensitivity of Table 2 of the main text, is shown beside them.

The molecular fraction of the bird tree has a range, whose ends follow the two rules of Materials and methods of the article (Four levels of coverage): 0.854 with every tip of the source's tree of phylogeny-placed taxa counted, and 0.843 without the tips whose only input is a taxonomy tree of the synthesis and those whose inputs could not be matched to the synthesis's list of inputs. Supplement S7 recomputes the levels at the lower end.

**Table S5 | Molecular tips of each reference tree.** Route: T, tips of a tree of the source's sequenced species; L, taxa of the source's alignment; E, per-tip evidence; S, every tip sequenced, as stated by the source. Mol. tips and Fraction: molecular tips and their share of the tips. Exact: molecular tips by exact label matching. Mol. species: distinct accepted species among the molecular tips. Source labels, Merged and Unmatched, for route T: the tip labels of the source's tree of sequenced species; those whose species key, after the names map, is that of another source label (subspecies of one species, or specimen labels the names map resolves to one name), which stand for one tip of the reference tree; and the source species absent from the reference tree (for Amphibians, Squamates and Nematodes, the outgroups removed from the reference tree, Supplement S2). Mol. tips = Source labels − Merged − Unmatched. Product: named species times the molecular fraction. [d]The source also distributes a dated tree of its molecular species. [g]The source also distributes a tree of its molecular species that is not dated: branch lengths in substitutions (Amphibians and Squamates) or no branch lengths (Birds).

| Group | Route | Tips | Mol. tips | Fraction | Exact | Source labels | Merged | Unmatched | Mol. species | Product |
|---|---|---|---|---|---|---|---|---|---|---|
| *Vertebrates* | | | | | | | | | | |
| Amphibians | T | 7,238 | 4,061[g] | 0.561 | 4,061 | 4,062 | 0 | 1 | 3,993 | 3897.7 |
| Birds | T | 10,824 | 9,239[g] | 0.854 | 9,239 | 9,239 | 0 | 0 | 8,980 | 8936.0 |
| Crocodilians | S | 25 | 25 | 1.000 | — | — | — | — | 20 | 20.0 |
| Fish | T | 31,347 | 11,596[d] | 0.370 | 11,561 | 11,638 | 42 | 0 | 11,510 | 11457.7 |
| Mammals | L | 5,911 | 4,098[d] | 0.693 | 0 | — | — | — | 3,945 | 3899.7 |
| Sharks | T | 1,192 | 610[d] | 0.512 | 610 | 610 | 0 | 0 | 603 | 588.5 |
| Squamates | T | 9,754 | 5,415[g] | 0.555 | 5,415 | 5,416 | 0 | 1 | 5,096 | 4970.3 |
| Turtles | S | 283 | 283 | 1.000 | — | — | — | — | 256 | 256.0 |
| *Plants* | | | | | | | | | | |
| Bryophytes | L | 533 | 533 | 1.000 | 532 | — | — | — | 505 | 505.0 |
| Ferns | L | 6,216 | 6,216 | 1.000 | 6,216 | — | — | — | 6,097 | 6097.0 |
| Seed plants | T | 348,886 | 76,688[d] | 0.220 | 76,341 | 79,874 | 3,186 | 0 | 70,723 | 61337.2 |
| *Arthropods* | | | | | | | | | | |
| Crustaceans | E | 95 | 95 | 1.000 | — | — | — | — | 65 | 65.0 |
| Insects | L | 53,373 | 53,373 | 1.000 | 53,373 | — | — | — | 42,078 | 42078.0 |
| Spiders | S | 1,431 | 1,431 | 1.000 | — | — | — | — | 888 | 888.0 |
| *Other animals* | | | | | | | | | | |
| Bivalves | L | 73 | 73 | 1.000 | 73 | — | — | — | 44 | 44.0 |
| Bryozoa | L | 565 | 565 | 1.000 | 565 | — | — | — | 337 | 337.0 |
| Cephalopods | L | 81 | 81 | 1.000 | 81 | — | — | — | 80 | 80.0 |
| Cnidaria | E | 289 | 289 | 1.000 | — | — | — | — | 258 | 258.0 |
| Echinoderms | E | 65 | 65 | 1.000 | — | — | — | — | 62 | 62.0 |
| Gastropods | L | 40 | 40 | 1.000 | 40 | — | — | — | 39 | 39.0 |
| Nematodes | T | 156 | 156 | 1.000 | 148 | 166 | 0 | 10 | 101 | 101.0 |
| Sponges | L | 119 | 119 | 1.000 | 111 | — | — | — | 86 | 86.0 |
| *Microbes & protists* | | | | | | | | | | |
| Diatoms | L | 1,113 | 1,113 | 1.000 | 0 | — | — | — | 39 | 39.0 |
| Fungi | L | 1,620 | 1,620 | 1.000 | 1,563 | — | — | — | 1,396 | 1396.0 |

### Supplement S5: time scale of the reference trees

A reference tree is ultrametric when the coefficient of variation of its root-to-tip distances, measured on the distributed file, is at most 0.05 (Table S6). The measured values fall into two separate sets: the largest among the 19 ultrametric trees is 0.0000 and the smallest among the others 0.2642, so any threshold between the two gives the same classification. An ultrametric tree is a chronogram in absolute time when its source states that its branch lengths are in millions of years set by a calibration, recorded with the evidence for each tree in the data release. The insect tree is ultrametric, its root height is below one and its source states no calibration, so it is in relative time and is not counted as dated. For each tree that is not a chronogram, the last column gives what its source study did, recorded with its evidence in the data release; for example, the one time tree archived with the fungal tree (Li et al., 2021) is a RelTime tree estimated "without fossil calibrations" (its STAR Methods), so the study dated the tree in relative time only.

**Table S6 | Time scale of the reference trees.** CV: coefficient of variation of the root-to-tip distances of the distributed file. Ultrametric: CV at most 0.05. Root height: in the units of the file (millions of years for a chronogram in absolute time). Dates: the codes of Table 1 of the main text (A, chronogram in absolute time; R, in relative time; S, the study dated the tree in relative time only; Dt, the study's archived chronogram lacks tips; Dn, the study dated the tree and did not archive the chronogram; Du, the archived chronogram could not be read; N, the study did not date the tree).

| Group | CV | Ultrametric | Root height | Dates |
|---|---|---|---|---|
| *Vertebrates* | | | | |
| Amphibians | 0.0000 | yes | 312.8 | A |
| Birds | 0.0000 | yes | 99.26 | A |
| Crocodilians | 0.0000 | yes | 87.14 | A |
| Fish | 0.0000 | yes | 368 | A |
| Mammals | 0.0000 | yes | 180.8 | A |
| Sharks | 0.0000 | yes | 365.4 | A |
| Squamates | 0.0000 | yes | 190 | A |
| Turtles | 0.0000 | yes | 212 | A |
| *Plants* | | | | |
| Bryophytes | 0.0000 | yes | 515 | A |
| Ferns | 0.0000 | yes | 421.8 | A |
| Seed plants | 0.0000 | yes | 325.1 | A |
| *Arthropods* | | | | |
| Crustaceans | 0.5887 | no | 0.9889 | Dt |
| Insects | 0.0000 | yes | 0.796 | R |
| Spiders | 0.0000 | yes | 366.1 | A |
| *Other animals* | | | | |
| Bivalves | 0.2642 | no | 14.55 | N |
| Bryozoa | 0.6176 | no | 4.564 | Dn |
| Cephalopods | 0.0000 | yes | 240.9 | A |
| Cnidaria | 0.0000 | yes | 573.1 | A |
| Echinoderms | 0.0000 | yes | 511.9 | A |
| Gastropods | 0.0000 | yes | 479.6 | A |
| Nematodes | 0.2973 | no | 1.158 | Du |
| Sponges | 0.0000 | yes | 526.4 | A |
| *Microbes & protists* | | | | |
| Diatoms | 0.0000 | yes | 190 | A |
| Fungi | 0.4018 | no | 4.672 | S |

### Supplement S6: crown ages and archived intervals

For each of the 18 reference chronograms, Table S7 gives the crown clade, its age and the interval archived for it. The crown clade is the clade the source study sampled, its ingroup. Where the source's outgroups belong to the group, they stay in the reference tree and are left out only when the crown node is found: the Scleractinia study samples antipatharian and corallimorpharian outgroups (Cnidaria), and the echinoid study samples crinoid, asteroid, ophiuroid and holothuroid outgroups (Echinoderms), so the crowns are those of Scleractinia and Echinoidea. The age is the source's own summary of the crown node: the annotated median height where the archive is an annotated summary tree, the median height over the set where it is a set of dated trees, and otherwise the height of the node in the archived chronogram; where the archive carries no interval, it is the root height of the reference tree. From a set of trees, the interval is the central 95% range of crown heights; from an annotated tree, it is the interval annotated on the node.

What an archive carries when date uncertainty is available, the last level of coverage, is defined in Materials and methods of the article (Four levels of coverage) and recorded per tree in the data release.

A set that adds taxonomically placed species to one dated tree does not carry it (t; Fish): pruned to its sequenced tips, each of the 100 trees of the fish set (Rabosky et al., 2018) has the branching times of the source's dated tree of sequenced species to within 0.001 million years, the width below which an interval counts as zero, so its crown age is the same in every tree (width 0.0 million years). A set whose dates are drawn from published studies onto one tree carries date uncertainty but no estimate of it (v; Birds). Table S7 also compares each crown interval with its calibrations; this is a description, and no level of coverage rests on it. The comparison excludes the crown interval in any of four cases. (z) Its width is below 0.001 million years (Bryophytes). (b) Its median lies within 0.01 million years of one of its ends (Diatoms). (p) It lies within the range that calibrations allow the crown and is not narrowed below half that range's width (Amphibians, Cephalopods, Gastropods, Mammals, Sponges, Squamates and Turtles): at least 90% of the interval lies inside the range, and the interval is at least 0.5 times as wide as the range. The range is that of a two-sided calibration the source places on the crown; for a crown without one, it is the band from the oldest minimum on the crown or on a node below it to the youngest maximum on the crown or on a node above it, where both exist (column Prior, marked [b]). (r) The set of trees was dated under root ages drawn from the calibration prior (Sharks), so its crown interval returns that prior by construction, whatever its containment and width ratio; the shark interval lies 82% inside the bounds of its prior, below the threshold of rule (p), and is excluded by rule (r). Rule (p) tests containment and width only. Where the calibration has a stated density, the column Quantiles gives the prior's cumulative probability at the interval's lower end, median and upper end, so a shift of the interval within the prior can be read: the mammal median lies at the prior's 0.84 quantile, the squamate interval excludes the upper part of a uniform prior (quantiles 0.03 to 0.73), and the turtle interval spans 0.57 of its prior's width. The prior's range is that of a run of the source's own input file sampling from the prior where one was run (Cephalopods; Materials and methods of the article, Four levels of coverage), otherwise the calibration as the source specifies it (recorded in the data release with the source's quotation for each calibration). A one-sided calibration, no calibration on the crown, or a calibration that could not be checked does not by itself exclude an interval, and a crown without a maximum on it or above it, or without a minimum on it or below it, has no range to compare with (column Prior, "no range"); the rules exclude intervals that match one of these patterns and do not show that the others were narrowed by the data. For a calibration with a stated density (a uniform, exponential, lognormal or gamma prior), the prior's range covers the same probability as the archived interval: its central range for an interval from a set of trees, its highest-density range for an annotated HPD interval. Two ranges in Table S7 are bounds that carry no stated probability: the shark root density is given by a hard minimum and a soft maximum, and the diatom crown by treePL hard bounds; neither outcome rests on them, since the shark set is excluded by rule (r) and the diatom median sits on a bound. The bands are bounds too: the amphibian band runs from the minimum of crown Batrachia to the upper end of the uniform root prior, the gastropod band from the soft minimum on node E (Sublitoidea), on or below the gastropod crown, to the soft maximum on the root, and the bryophyte band from the oldest minimum below the root to the root's maximum, on which the bryophyte interval sits. The interval is read from the set of dated trees that the summary tree summarises for Amphibians, Mammals, Squamates and Turtles; for Turtles the archived set is a sample of the posterior, so its median and interval differ slightly from those the source reports from the full posterior. The amphibian, mammal and squamate sets carry the posterior of a backbone dating (Materials and methods of the article, Four levels of coverage) (Jetz and Pyron, 2018; Upham et al., 2019; Tonini et al., 2016). The backbones are dated under node calibrations: for amphibians, fifteen uniform calibrations; for mammals, seventeen fossil calibrations with exponential priors and a root constraint; for squamates, secondary uniform calibrations from the posterior ranges of an earlier dated study. The amphibian crown carries no calibration of its own, but it lies between two two-sided calibrations: the root, one node up, has a uniform prior between the minimum and maximum ages of Amphibia–Amniota, and crown Batrachia, one node down, a minimum of 245.0 Ma and a maximum equal to the root's minimum age (Jetz and Pyron, 2018). The amphibian interval lies 100% inside the band from 245.0 to 350.1 Ma and fills 0.62 of its width, and the gastropod interval lies 100% inside the band from 418.0 to 550.0 Ma and fills 0.72 of it, so rule (p) excludes both. The two thresholds are conventions; Table S7 prints the containment and the width ratio of every interval with a two-sided calibration, so another threshold can be applied. The intervals of Cnidaria and Echinoderms are not excluded by any rule: the scleractinian crown has no range to compare with, since no maximum lies on it or on a node above it (Cnidaria), and the echinoid crown carries a two-sided calibration that its interval does not fill (Echinoderms: 81% inside the prior, width ratio 0.37, median at the prior's 0.91 quantile). No interval that the rules leave in lies on a crown whose calibration was not checked. No interval is archived for Crocodilians, Ferns, Seed plants and Spiders.

### Supplement S7: sensitivity analyses

Table S8 recomputes the four levels of coverage under alternatives to the denominator, to the set of trees and to the molecular fraction; some repeat columns of Table 2 of the main text for comparison.

*Denominators.* The insect count, the largest count of one group (Supplement S1), is lowered and raised by 20%. The Catalogue of Life count of a group is the number of accepted extant species under the taxa that define it in COL26.9. Where the base release holds fewer than 50% of the species that the extended release of the same version holds in a class, the extended count is used for that class (marked in Table S10). A second row uses base-release counts only and keeps the primary count of the 7 groups the base release does not cover.

*Groups and trees.* The denominator is restricted to the groups with a tree; and the seed-plant tree and the described seed plants are removed together, since that tree holds most of the named and dated species. The reference trees are classed by how they were assembled, with the classes of the figure of named coverage against the molecular fraction (Results of the article, Named against molecular coverage): 21 are primary analyses, 2 are compiled from published trees (Cephalopods and Crocodilians; the cephalopod study estimated branch lengths on the compiled topology from its own alignment, with a posterior set of dated trees), and 1 is an automated synthesis (Birds). Each class is removed in turn, and each is kept alone; the denominator stays the same, since removing a tree does not remove its species. The row "Named species of all trees" adds the named species of the other 237 eukaryotic trees of the catalogue, each assigned to a group by its Catalogue of Life family; its molecular value is the upper bound of Table 2 of the main text.

*Partition.* Table S9 counts the described eukaryotic species named in any tree of the catalogue eligible at a level (any tree) and, for a partition of these species into groups, the species each group's largest eligible tree names among them (one tree per group, the rule of the reference trees applied at the scale of the partition). The levels are read the same way on every tree: named, a tip that resolves to an accepted species of COL26.9; dated, in a tree dated in absolute time; date uncertainty available, in a reference tree whose archive carries it. They count named species, and the molecular level, enumerated only on the reference trees, is not compared. The groups of this paper assign a species outside the reference trees to the group whose reference trees carry its family most often, as the row "Named species of all trees" of Table S8 does; the other partitions use the COL26.9 classification of the species' family. One tree per group never counts more than any tree, and the two agree as the groups become smaller.

*Molecular fraction.* The bird row uses the lower end of the bird tree's molecular fraction (Supplement S4).

**Table S10 | Denominators and named coverage by group.** Described: the group's count and its source, used throughout the paper. CoL: the Catalogue of Life count; $^f$ a class holds fewer than 50% of the species of the extended release in the base release, and its extended count is used. Named: named coverage over the described count, over the Catalogue of Life count, and with the named species of all trees of the catalogue.

| | Described | | | Named (%) | | |
|---|---|---|---|---|---|---|
| Group | Species | Source | CoL | Described | CoL | All trees |
| *Vertebrates* | | | | | | |
| Fish | 36,288 | Eschmeyer's Catalog of Fishes (21 Sep. 2026) | 35,750$^f$ | 85.4 | 86.6 | 85.9 |
| Squamates | 12,252 | Reptile Database, June 2026 release (Uetz et al.) | 12,251 | 73.1 | 73.1 | 73.2 |
| Birds | 11,185 | IUCN Red List 2026-1, Table 1a (HBW-BirdLife v10) | 10,868 | 93.6 | 96.3 | 96.2 |

| Group | Species | Source | CoL | Described | CoL | All trees |
|---|---|---|---|---|---|---|
| Amphibians | 9,088 | AmphibiaWeb (11 Sep. 2026) | 8,916 | 76.4 | 77.9 | 85.9 |
| Mammals | 6,887 | Mammal Diversity Database v2.5 (2026) | 6,362 | 81.7 | 88.4 | 83.2 |
| Sharks | 1,331 | Eschmeyer's Catalog of Fishes (21 Sep. 2026) | 1,355[f] | 86.4 | 84.9 | 86.4 |
| Turtles | 359 | Turtle Taxonomy Working Group 2025 (10th ed.), living species | 370 | 71.3 | 69.2 | 76.0 |
| Hagfish | 93 | Eschmeyer's Catalog of Fishes (20 Oct. 2026) | 93 | — | — | — |
| Lampreys | 49 | Eschmeyer's Catalog of Fishes (21 Sep. 2026) | 49 | — | — | — |
| Crocodilians | 27 | The Reptile Database, June 2026 (via COL26.9) | 27 | 74.1 | 74.1 | 74.1 |
| Other vertebrates | 9 | The Reptile Database (Jun. 2026) and Eschmeyer's Catalog of Fishes (21 Sep. 2026) | 25[f] | — | — | — |
| *Plants* | | | | | | |
| Seed plants | 354,303 | World Flora Online, as in COL26.9 (2026-09-11) | 354,308[f] | 78.8 | 78.8 | 78.8 |
| Bryophytes | 19,273 | Catalogue of Life COL26.9 | 19,273 | 2.6 | 2.6 | 2.6 |
| Ferns | 11,999 | PPG II, pteridogroup/ppg v2.0.0 (15 Sep. 2026) | 12,457 | 50.8 | 48.9 | 50.8 |
| Green algae | 11,791 | AlgaeBase via Guiry 2024 (counts to Sep. 2023), extant | 4,912[f] | — | — | — |
| Rhodophyta | 7,276 | AlgaeBase via Guiry 2024 (1 Nov. 2023), living species | 3,546[f] | — | — | — |
| Lycophytes | 1,467 | PPG II, pteridogroup/ppg v2.0.0 (15 Sep. 2026) | 1,503 | — | — | — |
| *Arthropods* | | | | | | |
| Insects | 929,676 | Catalogue of Life COL26.9 | 929,676 | 4.5 | 4.5 | 4.8 |
| Crustaceans | 67,735 | Zhang 2013 (Zootaxa 3703) | 70,596[f] | 0.10 | 0.09 | 0.10 |
| Acari | 54,908 | Zhang 2013 (Zootaxa 3703), extant species | 28,874 | — | — | — |
| Spiders | 54,244 | World Spider Catalog v27 (read 4 Nov. 2026) | 53,379 | 1.6 | 1.7 | 1.6 |
| Myriapods | 16,072 | Chapman 2009 | 17,349[f] | — | — | — |
| Other arachnids | 12,765 | Catalogue of Life COL26.9 | 12,765 | — | — | — |
| Other hexapods | 10,138 | Catalogue of Life COL26.9 | 10,138[f] | — | — | — |
| Scorpiones | 2,954 | The Scorpion Files (Rein; Wayback 3 Oct. 2026) | 2,940 | — | — | — |
| Pycnogonida | 1,431 | WoRMS (Pycnobase) 2026-10-01 | 1,420 | — | — | — |
| *Other animals* | | | | | | |
| Gastropods | 76,288 | MolluscaBase, in Catalogue of Life COL26.9 | 76,288 | 0.05 | 0.05 | 0.05 |
| Flatworms | 29,487 | Zhang 2013 (29,488 species including 1 fossil) | 27,037 | — | — | — |
| Nematodes | 28,415 | Hodda 2022a (28,537 species including at least 122 fossil) | 19,598 | 0.36 | 0.52 | 0.36 |
| Annelids | 17,340 | Catalogue of Life COL26.9 | 17,340[f] | — | — | — |
| Cnidaria | 11,597 | WoRMS (accessed 4 Nov. 2026) | 11,582[f] | 2.2 | 2.2 | 2.2 |
| Bivalves | 10,043 | MolluscaBase, in Catalogue of Life COL26.9 | 10,043 | 0.44 | 0.44 | 0.44 |
| Sponges | 9,864 | World Porifera Database (15 Aug. 2026) | 9,873[f] | 0.87 | 0.87 | 0.87 |
| Echinoderms | 7,749 | WoRMS (accessed 4 Nov. 2026) | 7,737[f] | 0.80 | 0.80 | 0.80 |
| Bryozoa | 6,613 | WoRMS (accessed 4 Nov. 2026) | 5,418 | 5.1 | 6.2 | 5.1 |
| Other phyla | 3,863 | Catalogue of Life COL26.9 | 3,863[f] | — | — | — |
| Tunicates | 3,020 | Appeltans et al. 2012 | 3,146 | — | — | — |
| Myxozoa | 3,012 | World List of Myxozoa (WoRMS), in Catalogue of Life COL26.9 | 3,012 | — | — | — |
| Other Mollusca | 2,199 | MolluscaBase, in Catalogue of Life COL26.9 | 2,204[f] | — | — | — |
| Rotifera | 2,030 | Segers 2007 (about 2,030) | 2,471[f] | — | — | — |
| Tardigrada | 1,505 | Degma & Guidetti, Actual checklist of Tardigrada species (2025) | 1,451 | — | — | — |
| Acanthocephala | 1,325 | Catalogue of Life COL26.9 | 1,325 | — | — | — |
| Nemerteans | 1,275 | Kajihara et al. 2008 | 1,364 | — | — | — |
| Cephalopods | 868 | MolluscaBase, in Catalogue of Life COL26.9 | 868 | 9.2 | 9.2 | 9.2 |
| Ctenophores | 186 | WoRMS (accessed 4 Nov. 2026) | 182 | — | — | — |

| Group | Species | Source | CoL | Described | CoL | All trees |
|---|---|---|---|---|---|---|
| *Microbes & protists* | | | | | | |
| Fungi | 156,891 | Catalogue of Life COL26.9 (Species Fungorum Plus; Microsporidia) | 156,891 | 0.89 | 0.89 | 0.89 |
| Diatoms | 16,427 | AlgaeBase (Guiry 2024) | 6,723$^{f}$ | 0.24 | 0.58 | 0.24 |
| Foraminifera | 8,972 | World Foraminifera Database / WoRMS (4 Nov. 2026) | 8,928 | — | — | — |
| Apicomplexa | 6,000 | Morrison 2008 | 169$^{f}$ | — | — | — |
| Ciliates | 4,500 | Foissner et al. 2008 | 8,556$^{f}$ | — | — | — |
| Other protists | 3,353 | Pawlowski et al. 2012 (PLoS Biol 10: e1001419), Table S1: 29 lineages in no other group | 555$^{f}$ | — | — | — |
| Amoebozoa | 2,400 | Pawlowski et al. 2012 | 1,847$^{f}$ | — | — | — |
| Dinoflagellates | 2,377 | Gómez 2012 checklist (CICIMAR Oceánides 27: 65-140) | 1,425$^{f}$ | — | — | — |
| Brown algae | 2,094 | AlgaeBase (Guiry 2024) | 1,097$^{f}$ | — | — | — |
| Oomycetes | 1,673 | Catalogue of Life COL26.9 (Species Fungorum Plus) | 1,673 | — | — | — |
| Euglenozoa | 1,520 | CBOL Protist Working Group (Pawlowski et al. 2012), Table S1 | 578$^{f}$ | — | — | — |
| Chrysophyta | 1,216 | AlgaeBase (Guiry 2024) | 611$^{f}$ | — | — | — |
| Cercozoa | 600 | Pawlowski et al. 2012 (CBOL ProWG, Table S1) | 183$^{f}$ | — | — | — |
| Haptophytes | 517 | AlgaeBase (Guiry 2024; living species, 1 Nov. 2023) | 287$^{f}$ | — | — | — |
| Choanoflagellates | 244 | Carr et al. 2008 ('over 244') | 140$^{f}$ | — | — | — |

### Supplement S8: comparison with the Open Tree of Life

From the Open Tree of Life synthesis release v16.1 I took the leaves of the grafted solution, the tips placed by at least one input phylogeny, kept those that are extant species-rank taxa of the Open Tree taxonomy (OTT 3.7.3), and assigned each to the groups by its ancestry in that taxonomy. For each of the 21 groups whose molecular tips are listed, Table S11 gives the two comparisons of Materials and methods of the article (Comparison with the Open Tree of Life); the Open Tree names are resolved to accepted species by the rule of Supplement S3.

### Supplement S9: worked example

*Trees.* The trees fitted and their time axis are stated in Materials and methods of the article (Worked example); the crown ages to which they are not rescaled are those of Supplement S6. Rescaling by a constant $c$ adds the same constant, a function of $c$ and the number of tips, to every log-likelihood of the tree, so the likelihood-ratio statistics do not change. The pruned tree keeps the molecular tips of Supplement S4.

*Tree sets.* The tree sets are not redistributed: each is read from the source's archive at the address recorded in the data release, and the trees fitted are selected from it by index.

*Likelihood.* The model is the constant-rate birth–death process with each extant species sampled with probability $\rho$ (Stadler, 2009), conditioned on the crown age and on both crown lineages leaving sampled descendants. A node with more than two children is read as simultaneous splits, after internal branches of zero length are collapsed; Table S12 gives, for every fitted tree, its branching events and those at such nodes. The parameters are fitted as $r = \lambda - \mu > 0$ and $\varepsilon = \mu/\lambda$ in $[0, 1)$ by maximum likelihood in R 4.5.1.

*Intervals.* Each interval is a profile-likelihood interval: the values at which the log-likelihood lies within 1.92 of its maximum, half the 95% point of a chi-squared distribution with one degree of freedom. The same cutoff is used for every interval. The coverage is 95% only asymptotically and for interior values of $\varepsilon$; at $\varepsilon = 0$ the statistic is a mixture of a point mass at zero and a chi-squared distribution with one degree of freedom, so the interval is conservative there, and near zero its coverage is not uniform. The intervals exclude topological and dating uncertainty.

*Test of the full tree's fraction.* The restriction, the likelihood-ratio statistic, its reference distribution, the two conditions under which a test is reported and the family of tests are defined in Materials and methods of the article (Worked example). The two maximised log-likelihoods of each statistic are in Table S12, the

**Table S7 | Crown ages and archived intervals of the reference chronograms.** Age: crown age (millions of years). Interval and Width: the archived interval and its width (millions of years). Kind: post, posterior set of dated trees; boot, set of trees dated on bootstrap replicates; hpd, interval annotated on a summary tree; lab, interval in a node label; prior, root-age prior set: trees dated under root ages drawn from the calibration prior; tax, set that varies the placement of taxonomically added tips; date, set that varies the placement of taxonomically added tips and the dates drawn from published studies. Trees: trees in the set. Cal.: calibration on the crown (T, two-sided; O, one-sided; N, none; ?, not checked). Prior: range of a two-sided calibration on the crown ("no range" for an interval no rule excludes and that has nothing to be compared with): its central 95% range, its highest-density range for an annotated HPD interval, or, for Sharks and Diatoms, its bounds; [b]for a crown without one, the band from the oldest minimum on the crown or a node below it to the youngest maximum on the crown or a node above it. In prior: percentage of the interval inside the prior's range. Width ratio: interval width over prior width. Quantiles: cumulative probability of the calibration density as the source specifies it at the interval's lower end, median (the node height for an interval in a node label) and upper end; beyond a soft bound only the side is given. Parent: calibration on the node above the crown; Child: calibrations on the nodes below it, recorded for crowns without a two-sided calibration; both coded as Cal. Comparison: u, not excluded by any rule; t, no date uncertainty, the set varies only the placement of added tips; v, dates drawn from published studies; z, zero width; b, median at a bound; p, within the range calibrations allow the crown (Prior) and not narrowed below half its width; r, root ages drawn from the calibration prior; n, no interval archived. The comparison describes each interval against its calibrations as specified; how far the data moved an interval requires a run sampling from the prior, made for Cephalopods only.

| Group | Crown clade | Age | Interval | Width | Kind | Trees | Cal. | Prior | In prior | Width ratio | Quantiles | Parent | Child | Comparison |
|---|---|---|---|---|---|---|---|---|---|---|---|---|---|---|
| *Vertebrates* | | | | | | | | | | | | | | |
| Amphibians | Lissamphibia | 311.4 | 273.7–338.6 | 64.9 | post | 10,000 | N | 245.0–350.1[b] | 100 | 0.62 | — | T | T | p |
| Birds | Aves (crown birds) | 99.3 | 72.0–133.0 | 61 | date | 100 | — | — | — | — | — | — | — | v |
| Crocodilians | Crocodylia | 87.1 | — | — | — | — | — | — | — | — | — | — | — | n |
| Fish | Actinopterygii | 368.0 | 368.0–368.0 | 0 | tax | 100 | — | — | — | — | — | — | — | t |
| Mammals | Mammalia | 187.3 | 168.6–213.5 | 44.9 | post | 10,000 | T | 165.2–210.0 | 92 | 1.00 | 0.26, 0.84, 0.98 | — | — | p |
| Sharks | Chondrichthyes | 377.8 | 353.6–437.6 | 84 | prior | 10,000 | T | 333.6–422.4 | 82 | 0.95 | — | — | — | r |
| Squamates | Squamata | 186.4 | 177.0–203.1 | 26.1 | post | 10,000 | T | 176.9–212.3 | 100 | 0.74 | 0.03, 0.28, 0.73 | — | — | p |
| Turtles | Testudines | 205.4 | 179.3–231.4 | 52.1 | post | 100 | T | 158.0–249.0 | 100 | 0.57 | 0.25, 0.52, 0.79 | — | — | p |
| *Plants* | | | | | | | | | | | | | | |
| Bryophytes | Bryophyta s.l. (mosses + liverworts + hornworts) | 515.0 | 515.0–515.0 | 0 | boot | 1,000 | O | 383.0–515.0[b] | 0 | 0.00 | — | — | T | z |
| Ferns | Polypodiopsida | 421.8 | — | — | — | — | — | — | — | — | — | — | — | n |
| Seed plants | Spermatophyta | 325.1 | — | — | — | — | — | — | — | — | — | — | — | n |
| *Arthropods* | | | | | | | | | | | | | | |
| Spiders | Araneae | 366.1 | — | — | — | — | — | — | — | — | — | — | — | n |
| *Other animals* | | | | | | | | | | | | | | |
| Cephalopods | Coleoidea | 240.0 | 236.6–249.6 | 13 | post | 9,001 | T | 236.7–251.2 | 99 | 0.89 | 0.02, 0.39, 0.94 | — | — | p |
| Cnidaria | Scleractinia | 459.7 | 404.1–516.5 | 112 | hpd | 1 | N | no range | — | — | — | N | O | u |
| Echinoderms | Echinoidea | 294.4 | 280.2–303.2 | 23 | lab | 1 | T | 237.0–298.9 | 81 | 0.37 | 0.69, 0.91, >0.975 | T | — | u |
| Gastropods | Gastropoda | 479.6 | 434.2–528.8 | 94.6 | hpd | 1 | O | 418.0–550.0[b] | 100 | 0.72 | — | O | O | p |
| Sponges | Demospongiae | 526.4 | 514.6–540.8 | 26.2 | lab | 1 | T | 515.0–541.0 | 99 | 1.01 | <0.025, 0.44, 0.97 | — | — | p |
| *Microbes & protists* | | | | | | | | | | | | | | |
| Diatoms | Bacillariophyta | 190.0 | 190.0–228.8 | 38.8 | boot | 1,001 | T | 190.0–302.0 | 100 | 0.35 | 0.00, 0.00, 0.35 | — | — | b |

**Table S8 | Coverage at the four levels under alternatives.** Denominator: described species the percentages are formed over. Named, Molecular, Dated, Available (date uncertainty available): percentage of the denominator at each level. Last row: date uncertainty counted only where the archive holds dated trees of a posterior or bootstrap dating of sequence data whose crown age is neither fixed nor at its median on a hard bound (Amphibians, Cephalopods, Mammals, Squamates and Turtles). A dash marks a level the alternative does not define.

| Alternative | Denominator | Named | Molecular | Dated | Available |
|---|---|---|---|---|---|
| Reported | 2,059,063 | 19.2 | 7.6 | 5.5 | 1.2 |
| Insect count −20% | 1,873,128 | 21.1 | 8.4 | 6.0 | 1.3 |
| Insect count +20% | 2,244,998 | 17.6 | 7.0 | 5.0 | 1.1 |
| Catalogue of Life count for every group | 1,993,667 | 19.8 | 7.9 | 5.7 | 1.2 |
| Catalogue of Life count where the base release has one | 1,996,589 | 19.8 | 7.9 | 5.7 | 1.2 |
| Groups with a tree only | 1,839,402 | 21.5 | 8.5 | 6.2 | 1.3 |
| Seed plants removed (tree and described species) | 1,704,760 | 6.8 | 5.1 | 2.5 | 1.4 |
| Automated synthesis removed | 2,059,063 | 18.7 | 7.2 | 5.1 | 0.73 |
| Compiled trees removed | 2,059,063 | 19.2 | 7.6 | 5.5 | 1.2 |
| Primary analyses removed | 2,059,063 | 0.51 | 0.44 | 0.44 | 0.44 |
| Automated synthesis only | 2,059,063 | 0.51 | 0.44 | 0.44 | 0.44 |
| Compiled trees only | 2,059,063 | 0.00 | 0.00 | 0.00 | 0.00 |
| Primary analyses only | 2,059,063 | 18.7 | 7.2 | 5.1 | 0.72 |
| Bird molecular fraction at its lower end | 2,059,063 | 19.2 | 7.6 | 5.5 | 1.2 |
| Named species of all trees | 2,059,063 | 19.4 | ≤ 7.8 | — | — |
| Available: posterior or bootstrap dated sets with a free crown | 2,059,063 | — | — | — | 0.65 |

**Table S9 | Coverage with one tree per group, under five partitions.** Groups: groups that hold a named species. Any tree: described eukaryotic species named in any tree eligible at the level. One tree: the sum over groups of the species of the group named by its largest eligible tree. Ratio: one tree over any tree (%). Named, Dated, Available (date uncertainty available) as in the text.

| Partition | Groups | Level | Any tree | One tree | Ratio |
|---|---|---|---|---|---|
| groups of this paper | 24 | Named | 399,548 | 395,517 | 99.0 |
| | | Dated | 358,488 | 355,865 | 99.3 |
| | | Available | 34,469 | 34,469 | 100.0 |
| phyla | 26 | Named | 399,548 | 354,927 | 88.8 |
| | | Dated | 358,488 | 315,400 | 88.0 |
| | | Available | 34,469 | 11,501 | 33.4 |
| classes | 87 | Named | 399,548 | 395,160 | 98.9 |
| | | Dated | 358,488 | 355,564 | 99.2 |
| | | Available | 34,469 | 34,209 | 99.2 |
| orders | 535 | Named | 399,548 | 395,474 | 99.0 |
| | | Dated | 358,488 | 357,041 | 99.6 |
| | | Available | 34,469 | 34,421 | 99.9 |
| families | 3,290 | Named | 399,548 | 396,870 | 99.3 |
| | | Dated | 358,488 | 357,382 | 99.7 |
| | | Available | 34,469 | 34,466 | 100.0 |

**Table S11 | Molecular species of the reference trees and species the Open Tree of Life places by phylogeny, by group.** Open Tree: extant species-rank taxa placed by an input phylogeny in the group's clade. Over the 21 groups they sum to 128,032; the Open Tree's 129,889 eukaryotic species are these, 498 in the groups whose sequenced tips are not listed (Crocodilians, Spiders and Turtles), and 1,359 in the clade of no group. Accepted species: molecular species of the reference tree, those also among the Open Tree species after both are resolved to Catalogue of Life accepted species, those absent from them, and the percentage present. Exact names: distinct genus–epithet names of the sequenced tips and of the Open Tree species (an Open Tree name without a species key is left out, and names spelled alike count once, so the Open Tree column of this block is at most the taxon count), the names in both, and that number as a percentage of each.

| | | Accepted species | | | | Exact names | | | | |
|---|---|---|---|---|---|---|---|---|---|---|
| Group | Open Tree | Molecular | Both | Absent | % | Molecular | Open Tree | Both | % mol. | % OT |
| *Vertebrates* | | | | | | | | | | |
| Amphibians | 6,547 | 3,993 | 3,832 | 161 | 96.0 | 4,061 | 6,546 | 3,739 | 92.1 | 57.1 |
| Birds | 10,204 | 8,980 | 7,922 | 1,058 | 88.2 | 9,239 | 10,203 | 7,304 | 79.1 | 71.6 |
| Fish | 12,121 | 11,510 | 9,887 | 1,623 | 85.9 | 11,586 | 12,094 | 9,755 | 84.2 | 80.7 |
| Mammals | 3,801 | 3,945 | 2,771 | 1,174 | 70.2 | 4,098 | 3,769 | 2,707 | 66.1 | 71.8 |
| Sharks | 536 | 603 | 517 | 86 | 85.7 | 610 | 536 | 482 | 79.0 | 89.9 |
| Squamates | 4,136 | 5,096 | 3,754 | 1,342 | 73.7 | 5,415 | 4,136 | 3,829 | 70.7 | 92.6 |
| *Plants* | | | | | | | | | | |
| Bryophytes | 536 | 505 | 206 | 299 | 40.8 | 524 | 530 | 189 | 36.1 | 35.7 |
| Ferns | 3,835 | 6,097 | 3,462 | 2,635 | 56.8 | 6,216 | 3,818 | 3,104 | 49.9 | 81.3 |
| Seed plants | 54,671 | 70,723 | 32,730 | 37,993 | 46.3 | 75,692 | 54,475 | 33,580 | 44.4 | 61.6 |
| *Arthropods* | | | | | | | | | | |
| Crustaceans | 1,730 | 65 | 24 | 41 | 36.9 | 76 | 1,646 | 22 | 28.9 | 1.3 |
| Insects | 23,509 | 42,078 | 8,082 | 33,996 | 19.2 | 53,365 | 22,991 | 9,342 | 17.5 | 40.6 |
| *Other animals* | | | | | | | | | | |
| Bivalves | 29 | 44 | 0 | 44 | 0.0 | 50 | 29 | 0 | 0.0 | 0.0 |
| Bryozoa | 168 | 337 | 153 | 184 | 45.4 | 372 | 168 | 160 | 43.0 | 95.2 |
| Cephalopods | 284 | 80 | 41 | 39 | 51.2 | 81 | 242 | 41 | 50.6 | 16.9 |
| Cnidaria | 1,747 | 258 | 200 | 58 | 77.5 | 275 | 1,711 | 196 | 71.3 | 11.5 |
| Echinoderms | 45 | 62 | 13 | 49 | 21.0 | 64 | 45 | 13 | 20.3 | 28.9 |
| Gastropods | 332 | 39 | 2 | 37 | 5.1 | 39 | 332 | 1 | 2.6 | 0.3 |
| Nematodes | 196 | 101 | 43 | 58 | 42.6 | 131 | 182 | 54 | 41.2 | 29.7 |
| Sponges | 498 | 86 | 66 | 20 | 76.7 | 95 | 371 | 67 | 70.5 | 18.1 |
| *Microbes & protists* | | | | | | | | | | |
| Diatoms | 113 | 39 | 1 | 38 | 2.6 | 772 | 99 | 92 | 11.9 | 92.9 |
| Fungi | 2,994 | 1,396 | 294 | 1,102 | 21.1 | 1,495 | 2,981 | 258 | 17.3 | 8.7 |
| Total | 128,032 | 156,037 | 74,000 | 82,037 | 47.4 | 174,256 | 126,904 | 74,935 | 43.0 | 59.0 |

statistic over the trees of each archived set in Table S14, and its range under random resolution of the polytomies of the consensus trees in Table S15. The second condition, node ages estimated by a dating analysis, is not met by the bird summary tree: 3,836 of 10,823 internal nodes (35.4%), and 2,251 of 9,238 on the tree pruned to its sequenced tips (24.4%), carry no published age, their ages interpolated or set at taxon addition (measured from the source's published ages per node and recorded in the data release); before taxon addition its tree of the 9,239 tips placed by input phylogenies has 231 branching events at polytomies, so the binary pruned tree carries 118 nodes that no input phylogeny resolves. A group whose summary-tree statistic is zero stays in the family of tests.

*Sampling by family.* In the family fit each tip carries the sampling fraction of its Catalogue of Life family: the family's accepted species among the sequenced tips over its accepted species in COL26.9, at most one; a tip without a resolved family carries the tree's uniform fraction. The likelihood is the approximation described in Materials and methods of the article (Worked example); at a node whose children carry different values it takes the value of the first child stored. Taking the value of the last child instead changes the log-likelihood at the same parameters by up to 12.6 units and the estimates by up to 0.1% in $r$ and 0.0021 in $\varepsilon$ (Table S13). With every fraction equal, the likelihood returns the uniform one: on every tree the two log-likelihoods differ by less than $10^{-6}$. The family fractions pooled over families (accepted molecular species over Catalogue species) differ from the uniform fraction, which divides molecular tips by the group's described count (Table S13). The family fractions are in the data release.

*Archived tree sets.* The fits over the dated trees a source archives follow Materials and methods of the article (Worked example); the sets, the sample and the fits are in the data release, and Table S14 gives the fits per set. The sequenced tips of the reference tree are matched in each tree of a posterior or date-assignment set by their source labels. A dated tree that a source archives of its sequenced species alone is fitted whole with the fraction its tips give. The shark taxon-complete trees, pruned to the sequenced species, are not the source's trees of those species: their topology differs at a median of 129 of 609 clades, a median of 307 clades have ages within 1% of the closest tree of the sequenced-species set, the root ages differ by a median of 0.231 Myr, and they carry more nodes younger than 5 Myr (median 51 against 38). The summary-tree fits of the groups for which Table 3 of the main text and Fig. 6 of the main text give the median and range over a set are in Table S12. On the full consensus trees of Amphibians and Squamates the profile intervals of $r$ and $\varepsilon$, like the estimates, lie outside the range over every tree of the posterior set.

*Polytomies.* As a check of the consensus summaries only, every uniform-sampling fit was repeated 3 times with the polytomies resolved at random: of the $k-1$ events of a node with $k$ children, one stays at the node's age and the others are drawn uniformly between that age and the age of the node's oldest child. Under uniform sampling the likelihood depends on the branching times only, so no topology is drawn. Table S15 gives the estimates as fitted and their range over the resolutions. The resolutions use one placement rule and few draws, so their range does not describe the dating uncertainty of these trees; the archived tree sets above do.

*Checks.* The fitted rates agree with phytools::fit.bd on the 17 uniform-sampling fits of trees with at most 15,000 tips (the full fish and seed-plant trees and the pruned seed-plant tree, fitted with two fractions, are larger), to within 0.01% (difference of $\lambda$ and of $\mu$ relative to $\lambda$: where $\mu$ is small, estimates that differ by more than 0.01% of $\mu$ can have log-likelihoods equal to six decimals). The acceptance test of the data release recomputes the log-likelihood of every uniform-sampling fit of up to that size with a separate implementation, and checks that the profile statistic at each interval end equals the cutoff.

### Supplement S10: data and code

The archive is data release v1.1.5 of the repository named under Data availability. It holds the redistributed trees of the catalogue, with their tips labelled by identifiers and a dictionary of those labels; the per-group table; the lists of molecular tips; the resolution of the tip labels to accepted species; the registers of every decision taken for a tree, each with its evidence; the generated numbers, tables and figure data of the article and of this Supplement; the code of the scheduled procedure that keeps the catalogue current (Materials and methods of the article); the code that rebuilds every derived file from the trees and the registers, with the scripts that produce the inputs needing a network, R 4.5.1 or data that are not redistributed; and a script

**Table S12 | Every birth–death fit of the worked example.** Fit: tree (full, or pruned to its molecular tips) and sampling fraction (own: tips over described species; full-tree: the full tree's fraction; family: one fraction per family). $\lambda$, $\mu$: speciation and extinction rates (per million years); $r$ and $\varepsilon$ with 95% profile-likelihood intervals; log-likelihood at the maximum. Events: branching events of the fitted tree; at polytomies: those at nodes with more than two children.

| Group | Fit | Tips | $\rho$ | $\lambda$ | $\mu$ | $r$ | $\varepsilon$ | Log-likelihood | Events | At polytomies |
|---|---|---|---|---|---|---|---|---|---|---|
| Amphibians | full, own $\rho$ | 7,238 | 0.80 | 0.0338 | 0 | 0.0338 (0.0330–0.0345) | 0.000 (0.000–0.004) | −32500.86 | 7,237 | 3,513 |
| | pruned, own $\rho$ | 4,061 | 0.45 | 0.0624 | 0.0216 | 0.0407 (0.0385–0.0430) | 0.347 (0.276–0.411) | −17769.59 | 4,060 | 1,176 |
| | pruned, full-tree $\rho$ | 4,061 | 0.80 | 0.038 | 0 | 0.0380 (0.0369–0.0391) | 0.000 (0.000–0.032) | −17773.67 | 4,060 | 1,176 |
| | pruned, family $\rho$ | 4,061 | — | 0.0564 | 0.0137 | 0.0426 (0.0404–0.0448) | 0.244 (0.163–0.318) | −17659.04 | 4,060 | 1,176 |
| Birds | full, own $\rho$ | 10,824 | 0.97 | 0.148 | 0 | 0.1478 (0.1425–0.1506) | 0.000 (0.000–0.058) | −31696.70 | 10,823 | 0 |
| | pruned, own $\rho$ | 9,239 | 0.83 | 0.158 | 0.0118 | 0.1463 (0.1403–0.1522) | 0.075 (0.008–0.138) | −27531.50 | 9,238 | 0 |
| | pruned, full-tree $\rho$ | 9,239 | 0.97 | 0.14 | 0 | 0.1403 (0.1373–0.1432) | 0.000 (0.000–0.027) | −27534.00 | 9,238 | 0 |
| | pruned, family $\rho$ | 9,239 | — | 0.156 | 0.00761 | 0.1487 (0.1428–0.1545) | 0.049 (0.000–0.113) | −27433.66 | 9,238 | 0 |
| Fish | full, own $\rho$ | 31,347 | 0.86 | 0.216 | 0.172 | 0.0434 (0.0417–0.0451) | 0.799 (0.789–0.809) | −101696.35 | 31,346 | 0 |
| | pruned, own $\rho$ | 11,596 | 0.32 | 0.19 | 0.142 | 0.0480 (0.0461–0.0499) | 0.748 (0.731–0.764) | −45366.45 | 11,595 | 0 |
| | pruned, full-tree $\rho$ | 11,596 | 0.86 | 0.0703 | 0.0223 | 0.0480 (0.0461–0.0499) | 0.318 (0.272–0.361) | −45366.45 | 11,595 | 0 |
| | pruned, family $\rho$ | 11,596 | — | 0.152 | 0.0992 | 0.0524 (0.0505–0.0544) | 0.654 (0.632–0.676) | −45269.30 | 11,595 | 0 |
| Mammals | full, own $\rho$ | 5,911 | 0.86 | 0.298 | 0.214 | 0.0846 (0.0778–0.0915) | 0.716 (0.685–0.746) | −16486.65 | 5,910 | 0 |
| | pruned, own $\rho$ | 4,098 | 0.60 | 0.29 | 0.202 | 0.0884 (0.0812–0.0958) | 0.695 (0.658–0.730) | −12419.84 | 4,097 | 0 |
| | pruned, full-tree $\rho$ | 4,098 | 0.86 | 0.201 | 0.113 | 0.0884 (0.0812–0.0958) | 0.560 (0.506–0.610) | −12419.84 | 4,097 | 0 |
| | pruned, family $\rho$ | 4,098 | — | 0.275 | 0.185 | 0.0895 (0.0823–0.0969) | 0.674 (0.634–0.711) | −12432.28 | 4,097 | 0 |
| Seed plants | full, own $\rho$ | 348,886 | 0.98 | 0.0888 | 0.0276 | 0.0612 (0.0607–0.0617) | 0.311 (0.303–0.319) | −1256081.65 | 348,885 | 275,542 |
| | pruned, own $\rho$ | 76,688 | 0.22 | 1.67 | 1.62 | 0.0469 (0.0454–0.0484) | 0.972 (0.971–0.973) | −206943.71 | 76,687 | 124 |
| | pruned, full-tree $\rho$ | 76,688 | 0.98 | 0.367 | 0.32 | 0.0469 (0.0454–0.0484) | 0.872 (0.867–0.877) | −206943.71 | 76,687 | 124 |
| | pruned, family $\rho$ | 76,688 | — | 1.58 | 1.53 | 0.0498 (0.0482–0.0513) | 0.968 (0.967–0.970) | −209157.25 | 76,687 | 124 |
| Sharks | full, own $\rho$ | 1,192 | 0.90 | 0.0487 | 0.028 | 0.0207 (0.0176–0.0239) | 0.575 (0.474–0.661) | −5301.16 | 1,191 | 0 |
| | pruned, own $\rho$ | 610 | 0.46 | 0.0466 | 0.0252 | 0.0214 (0.0180–0.0249) | 0.541 (0.398–0.655) | −2945.74 | 609 | 0 |
| | pruned, full-tree $\rho$ | 610 | 0.90 | 0.0239 | 0.00246 | 0.0214 (0.0180–0.0243) | 0.103 (0.000–0.325) | −2945.74 | 609 | 0 |
| | pruned, family $\rho$ | 610 | — | 0.0439 | 0.022 | 0.0219 (0.0185–0.0255) | 0.501 (0.347–0.624) | −2954.61 | 609 | 0 |
| Squamates | full, own $\rho$ | 9,754 | 0.80 | 0.0569 | 0 | 0.0569 (0.0558–0.0580) | 0.000 (0.000–0.004) | −38743.75 | 9,753 | 4,362 |
| | pruned, own $\rho$ | 5,415 | 0.44 | 0.0826 | 0.0127 | 0.0699 (0.0669–0.0730) | 0.153 (0.076–0.225) | −21412.38 | 5,414 | 1,319 |
| | pruned, full-tree $\rho$ | 5,415 | 0.80 | 0.0576 | 0 | 0.0576 (0.0562–0.0590) | 0.000 (0.000–0.010) | −21454.25 | 5,414 | 1,319 |
| | pruned, family $\rho$ | 5,415 | — | 0.0815 | 0.0107 | 0.0708 (0.0677–0.0738) | 0.131 (0.052–0.205) | −21478.75 | 5,414 | 1,319 |

**Table S13 | Family sampling fractions of the pruned trees.** Families: Catalogue of Life families among the molecular tips. Tips with a family: molecular tips whose family fraction is defined, of all molecular tips. Fraction: minimum, median and maximum over families, and pooled over families (accepted molecular species over Catalogue species). Uniform $\rho$: the fraction of the uniform fit (molecular tips over described species). Child order: $\varepsilon$ and $r$ of the family fit with the value at a node taken from the first and from the last child.

| | | Tips | | Fraction | | | | | Child order | | | |
|---|---|---|---|---|---|---|---|---|---|---|---|---|
| Group | Families | With a family | All | Min. | Median | Max. | Pooled | Uniform $\rho$ | $\varepsilon$, first | $\varepsilon$, last | $r$, first | $r$, last |
| Amphibians | 78 | 4,054 | 4,061 | 0.08 | 0.50 | 1.00 | 0.45 | 0.45 | 0.2438 | 0.2435 | 0.04261 | 0.04263 |
| Birds | 254 | 9,004 | 9,239 | 0.24 | 0.94 | 1.00 | 0.81 | 0.83 | 0.0487 | 0.0488 | 0.14872 | 0.14871 |
| Fish | 496 | 11,550 | 11,596 | 0.02 | 0.41 | 1.00 | 0.32 | 0.32 | 0.6543 | 0.6542 | 0.05241 | 0.05240 |
| Mammals | 159 | 3,989 | 4,098 | 0.14 | 0.73 | 1.00 | 0.61 | 0.60 | 0.6739 | 0.6736 | 0.08954 | 0.08956 |
| Seed plants | 450 | 73,858 | 76,688 | 0.01 | 0.34 | 1.00 | 0.20 | 0.22 | 0.9684 | 0.9684 | 0.04976 | 0.04976 |
| Sharks | 63 | 610 | 610 | 0.06 | 0.50 | 1.00 | 0.45 | 0.46 | 0.5010 | 0.5031 | 0.02192 | 0.02190 |
| Squamates | 75 | 5,125 | 5,415 | 0.02 | 0.50 | 1.00 | 0.42 | 0.44 | 0.1314 | 0.1307 | 0.07077 | 0.07081 |

**Table S14 | Fits over the dated trees the sources archive.** Set: the source's posterior set; its sets with dates drawn at random or as means of published dates (Birds); its root-age prior set, dated under root ages drawn from the calibration prior (Sharks); or its dated trees of its sequenced species alone. Trees: trees fitted. Tips: of the pruned tree and of the full tree (outgroups removed). $\varepsilon$, matching: median (range) of the extinction fraction with the matching fraction. LR: median (range) of the likelihood-ratio statistic of the full tree's fraction on the pruned tree. Binds: trees in which the full tree's fraction forces extinction to zero, a property of each tree. $r$ ratio: range of $r$ on the full over $r$ on the pruned tree. Summary tree: $\varepsilon$, matching, and LR of the reference tree (Table S12). The mammal trees of sequenced species include *Nyctinomops kalinowskii*, which the source's taxonomy table, the list of sequenced tips used for the reference tree, marks as not sampled, so their tips exceed the reference tree's sequenced tips by that species; the mammal tree sets hold no outgroup, and the outgroups of the amphibian and squamate sets are removed before fitting.

| | | | Tips | | | | | | Summary tree | |
|---|---|---|---|---|---|---|---|---|---|---|
| Group | Set | Trees | Pruned | Full | $\varepsilon$, matching | LR | Binds | $r$ ratio | $\varepsilon$ | LR |
| Amphibians | posterior set | 100 | 4,061 | 7,238 | 0.40 (0.28–0.48) | 1.80 (0.00–20.53) | 89 | 0.97–1.05 | 0.35 | 8.16 |
| Birds | date assignment, random dates | 100 | 9,239 | 10,824 | 0.20 (0.10–0.27) | 0.00 (0.00–2.35) | 2 | 1.00–1.02 | 0.07 | 4.99 |
| | date assignment, mean dates | 100 | 9,239 | 10,824 | 0.19 (0.18–0.19) | 0.00 (0.00) | 0 | 0.99–1.01 | 0.07 | 4.99 |
| Mammals | posterior set | 100 | 4,098 | 5,911 | 0.68 (0.66–0.71) | 0.00 (0.00) | 0 | 0.92–0.98 | 0.70 | 0.00 |
| | source's tree of sequenced species | 100 | 4,099 | — | 0.71 (0.68–0.74) | 0.00 (0.00) | 0 | — | 0.70 | 0.00 |
| Sharks | root ages drawn from the prior | 100 | 610 | 1,192 | 0.54 (0.31–0.72) | 0.00 (0.00–4.40) | 7 | 0.89–1.07 | 0.54 | 0.00 |
| | source's tree of sequenced species | 100 | 610 | — | 0.43 (0.09–0.64) | 0.68 (0.00–16.76) | 84 | — | 0.54 | 0.00 |
| Squamates | posterior set | 100 | 5,415 | 9,754 | 0.27 (0.14–0.41) | 35.57 (2.03–90.73) | 100 | 0.92–0.99 | 0.15 | 83.74 |

**Table S15 | Fits with the polytomies resolved at random.** Events: branching events at polytomies, and all branching events, of the full and the pruned tree. $r$ ratio: $r$ on the full over $r$ on the pruned tree, each with its own fraction. $\varepsilon$, matching: extinction fraction of the pruned tree with the matching fraction. LR: likelihood-ratio statistic of the full tree's fraction on the pruned tree. Each quantity as fitted, and its range over the 3 random resolutions. Outcome of the full tree's fraction on the pruned tree under every resolution: rescaled, or $\mu = 0$; no test is made on these trees, whose summary trees carry polytomies.

| | Events, full | | Events, pruned | | $r$ ratio | | $\varepsilon$, matching | | LR | | |
|---|---|---|---|---|---|---|---|---|---|---|---|
| Group | Polytomy | All | Polytomy | All | Fitted | Resolved | Fitted | Resolved | Fitted | Resolved | Outcome |
| Amphibians | 3,513 | 7,237 | 1,176 | 4,060 | 0.83 | 0.89–0.90 | 0.35 | 0.32–0.34 | 8.16 | 9.50–12.45 | $\mu = 0$ |
| Birds | 0 | 10,823 | 0 | 9,238 | 1.01 | 1.01 | 0.07 | 0.07 | 4.99 | 4.99 | $\mu = 0$ |
| Fish | 0 | 31,346 | 0 | 11,595 | 0.90 | 0.90 | 0.75 | 0.75 | 0.00 | 0.00 | rescaled |
| Mammals | 0 | 5,910 | 0 | 4,097 | 0.96 | 0.96 | 0.70 | 0.70 | 0.00 | 0.00 | rescaled |
| Seed plants | 275,542 | 348,885 | 124 | 76,687 | 1.30 | 1.23 | 0.97 | 0.97 | 0.00 | 0.00 | rescaled |
| Sharks | 0 | 1,191 | 0 | 609 | 0.97 | 0.97 | 0.54 | 0.54 | 0.00 | 0.00 | rescaled |
| Squamates | 4,362 | 9,753 | 1,319 | 5,414 | 0.81 | 0.91–0.92 | 0.15 | 0.20–0.21 | 83.74 | 58.76–63.02 | $\mu = 0$ |

that recomputes every number printed in the article and in this Supplement, the cells of their tables and the data of the figures, from the archived files alone, using only the Python standard library, and compares each with the generated files. Each tree file is redistributed under the licence of its source, which the release states file by file. For each of the 8 trees that are not redistributed (named in the caption of Table S1), the release gives the source to download it from, the checksum, tip count and tip labels of the file used here and how it was derived; every number of the article and of this Supplement is counted on all 263 trees.

*Branch lengths of the distributed trees.* Each tree of release v1.1.5 is checked against the file it is built from. The reference trees, 26 in all with those of Bacteria and Archaea, are their archived sources with the removed and merged tips taken out (Supplement S2); the branch lengths of each archived source equal those of the source's own file (Supplement S1). Every other tree equals, token for token after the tip labels are mapped, a copy made before tips were relabelled with dictionary identifiers. In data releases v1.0.3 to v1.0.8, that relabelling also rewrote digits that are not tip labels, so the files of 256 of these 263 trees carry altered branch lengths (854,760 branches) and altered numeric node labels, such as support values and stored age intervals; the relabelling did not change tip sets or topologies. Analyses that used branch lengths, node ages or node labels from those releases should be repeated with the trees of release v1.1.5. The erratum of the release lists the releases and the files.

Orr, R. J. S., Di Martino, E., Ramsfjell, M. H., Gordon, D. P., Berning, B., Chowdhury, I., Craig, S., Cumming, R. L., Figuerola, B., Florence, W., Harmelin, J.-G., Hirose, M., Huang, D., Jain, S. S., Jenkins, H. L., Kotenko, O. N., Kuklinski, P., Lee, H. E., Madurell, T., McCann, L., Mello, H. L., Obst, M., Ostrovsky, A. N., Paulay, G., Porter, J. S., Shunatova, N. N., Smith, A. M., Souto-Derungs, J., Vieira, L. M., Voje, K. L., Waeschenbach, A.,

Zágoršek, K., Warnock, R. C. M. and Liow, L. H. (2022), 'Paleozoic origins of cheilostome bryozoans and their parental care inferred by a new genome-skimmed phylogeny', *Science Advances* **8**(13), eabm7452.

Parks, D. H., Chaumeil, P.-A., Mussig, A. J., Rinke, C., Chuvochina, M. and Hugenholtz, P. (2026), 'GTDB release 10: a complete and systematic taxonomy for 715,230 bacterial and 17,245 archaeal genomes', *Nucleic Acids Research* **54**(D1), D743–D754.

Pfeiffer, J. M., Breinholt, J. W. and Page, L. M. (2019), 'Unioverse: a phylogenomic resource for reconstructing the evolution of freshwater mussels (Bivalvia, Unionoida)', *Molecular Phylogenetics and Evolution* **137**, 114–126.

Portik, D. M., Streicher, J. W. and Wiens, J. J. (2023), 'Frog phylogeny: a time-calibrated, species-level tree based on hundreds of loci and 5,242 species', *Molecular Phylogenetics and Evolution* **188**, 107907.

Qing, X., Zhang, Y. M., Sun, S., Ahmed, M., Lo, W.-S., Bert, W., Holovachov, O. and Li, H. (2025), 'Phylogenomic insights into the evolution and origin of Nematoda', *Systematic Biology* **74**(3), 349–358.

Rabosky, D. L., Chang, J., Title, P. O., Cowman, P. F., Sallan, L., Friedman, M., Kaschner, K., Garilao, C., Near, T. J., Coll, M. and Alfaro, M. E. (2018), 'An inverse latitudinal gradient in speciation rate for marine fishes', *Nature* **559**(7714), 392–395.

Ran, J.-H., Shen, T.-T., Wu, H., Gong, X. and Wang, X.-Q. (2018), 'Phylogeny and evolutionary history of Pinaceae updated by transcriptomic analysis', *Molecular Phylogenetics and Evolution* **129**, 106–116.

Romiguier, J., Borowiec, M. L., Weyna, A., Helleu, Q., Loire, E., La Mendola, C., Rabeling, C., Fisher, B. L., Ward, P. S. and Keller, L. (2022), 'Ant phylogenomics reveals a natural selection hotspot preceding the origin of complex eusociality', *Current Biology* **32**(13), 2942–2947.e4.

Särkinen, T., Bohs, L., Olmstead, R. G. and Knapp, S. (2013), 'A phylogenetic framework for evolutionary study of the nightshades (Solanaceae): a dated 1000-tip tree', *BMC Evolutionary Biology* **13**(1), 214.

Smith, S. A. and Brown, J. W. (2018), 'Constructing a broadly inclusive seed plant phylogeny', *American Journal of Botany* **105**(3), 302–314.

Stadler, T. (2009), 'On incomplete sampling under birth–death models and connections to the sampling-based coalescent', *Journal of Theoretical Biology* **261**(1), 58–66.

Stein, R. W., Mull, C. G., Kuhn, T. S., Aschliman, N. C., Davidson, L. N. K., Joy, J. B., Smith, G. J., Dulvy, N. K. and Mooers, A. O. (2018), 'Global priorities for conserving the evolutionary history of sharks, rays and chimaeras', *Nature Ecology & Evolution* **2**(2), 288–298.

Stewart, A. A. and Wiens, J. J. (2025), 'A time-calibrated salamander phylogeny including 765 species and 503 genes', *Molecular Phylogenetics and Evolution* **204**, 108272.

Tagliacollo, V. A., Tan, M., Reis, R. E., Gaia, R., Carrijo, V., Ranuzi, M., Craig, J. M. and Albert, J. S. (2024), 'Time-calibrated phylogeny of Neotropical freshwater fishes', *Frontiers in Bioinformatics* **4**, 1433995.

Thompson, J. B., Davis, K. E., Dodd, H. O., Wills, M. A. and Priest, N. K. (2023), 'Speciation across the Earth driven by global cooling in terrestrial orchids', *Proceedings of the National Academy of Sciences* **120**(29), e2102408120.

Thompson, J. B., Martinez, C., Avaria-Llautureo, J., Ramírez-Barahona, S., Manzanarez-Villasana, G., Culham, A., Gdaniec, A., Ryan, G., Venditti, C., Keeling, G. and Priest, N. K. (2026), 'CactEcoDB: trait, spatial, environmental, phylogenetic and diversification data for the cactus family', *Scientific Data* **13**(1), 623.

Thomson, R. C., Spinks, P. Q. and Shaffer, H. B. (2021), 'A global phylogeny of turtles reveals a burst of climate-associated diversification on continental margins', *Proceedings of the National Academy of Sciences* **118**(7), e2012215118.

Tonini, J. F. R., Beard, K. H., Ferreira, R. B., Jetz, W. and Pyron, R. A. (2016), 'Fully-sampled phylogenies of squamates reveal evolutionary patterns in threat status', *Biological Conservation* **204**, 23–31.

Upham, N. S., Esselstyn, J. A. and Jetz, W. (2019), 'Inferring the mammal tree: species-level sets of phylogenies for questions in ecology, evolution, and conservation', *PLOS Biology* **17**(12), e3000494.

Vaga, C. F., Quattrini, A. M., Galvão de Lossio e Seiblitz, I., Huang, D., Quek, Z. B. R., Stolarski, J., Cairns, S. D. and Kitahara, M. V. (2025), 'A global coral phylogeny reveals resilience and vulnerability through deep time', *Nature* **648**(8093), 377–382.

Williamson, K., Eme, L., Baños, H., McCarthy, C. G. P., Susko, E., Kamikawa, R., Orr, R. J. S., Muñoz-Gómez, S. A., Minh, B. Q., Simpson, A. G. B. and Roger, A. J. (2025), 'A robustly rooted tree of eukaryotes reveals their excavate ancestry', *Nature* **640**(8060), 974–981.

Wolfe, J. M., Breinholt, J. W., Crandall, K. A., Lemmon, A. R., Lemmon, E. M., Timm, L. E., Siddall, M. E. and Bracken-Grissom, H. D. (2019), 'A phylogenomic framework, evolutionary timeline and genomic resources for comparative studies of decapod crustaceans', *Proceedings of the Royal Society B* **286**(1901), 20190079.

Zapata, F., Wilson, N. G., Howison, M., Andrade, S. C. S., Jörger, K. M., Schrödl, M., Goetz, F. E., Giribet, G. and Dunn, C. W. (2014), 'Phylogenomic analyses of deep gastropod relationships reject Orthogastropoda', *Proceedings of the Royal Society B* **281**(1794), 20141739.
